# A pathway to produce non-coding piRNAs from endogenous protein-coding regions supports Drosophila spermatogenesis

**DOI:** 10.1101/2022.10.02.510544

**Authors:** Taichiro Iki, Shinichi Kawaguchi, Toshie Kai

**Author notes:** Correspondence Lead contact: Taichiro Iki.

## Abstract

PIWI-interacting (pi)RNA pathways control transposable elements (TEs) and endogenous genes in animal gonads, playing important roles in gamete formation. Here, we report a mechanism by which <u>endo</u>genous protein-coding regions, that normally provide their sequences for translation, serve as origins of non-coding piRNA biogenesis in *Drosophila melanogaster* testes. The products, namely endo-piRNAs, formed silencing complexes with Aubergine (Aub) in germ cells. Proximity proteome combined to functional analyses revealed a testis-specialized chaperone, Cyclophilin 40 (Cyp40), selectively increases endo-piRNA occupancy inside Aub-RISCs aside from other TE-related piRNAs. Moreover, Argonaute 2 (Ago2) activities were found critical for endo-piRNA production. We provide evidence that Ago2-bound short interfering (si)RNAs and micro(mi)RNAs specify precursors and direct endo-piRNA biogenesis. Consistently, Aub and Ago2 cooperate in spermatid differentiation and regulate endogenous genes via endo-piRNA-directed mRNA cleavage. Collectively, our data highlight that Drosophila testes employ a unique strategy to expand the diversity of germline piRNAs supporting late spermatogenesis.

**Headlines:**

- Endogenous protein-coding regions derive non-coding endo-piRNAs
- endo-piRNA and TE-piRNA are produced via distinct mechanisms
- siRNA and miRNA activities direct secondary piRNA biogenesis
- endo-piRNA pathway controls chromatin and sperm formation

## Introduction

In eukaryotes, diverse biological processes are controlled by silencing mechanisms relying on 20∼30-nucleotide (nt) small non-coding RNAs. Small RNAs interact with Argonaute (Ago) family proteins inside RNA-induced silencing complexes (RISCs), acting as sequence-dependent guide for selecting silencing targets. PIWI-interacting (pi)RNAs comprise a group of small RNAs forming RISCs with PIWI-clade Ago members and accumulating in animal gonads. piRNA-directed mechanisms are crucial in defending germline genomes against invasions of transposable elements (TEs) (Czech et al., 2018). PIWI proteins and piRNAs also play key roles in regulating endogenous genes (Ramat and Simonelig, 2021). Hence, defective piRNA pathways can lead to failure in stem cell maintenance, functional gamete formation, and other processes. However, the diversity, biogenesis, and functions of piRNAs are not fully elucidated.

Selected transcripts can serve as precursors for piRNA production. The major sources of precursors are defined intergenic regions called piRNA clusters, where truncated TEs can accumulate as remnant of past invasive activities (Aravin et al., 2007; Brennecke et al., 2007). In *Drosophila melanogaster*, cluster regions are largely heterochromatinized, and thus require a non-canonical mechanism for transcription initiation by RNA polymerase II. In germ cells, this relies on complexes containing heterochromatin protein 1 variant, Rhino (Rhi) and the cofactors, Deadlock and Cutoff (Andersen et al., 2017; Chen et al., 2016; Czech et al., 2013; Klattenhoff et al., 2009; Mohn et al., 2014). Distinguished from canonical transcripts including mRNAs, cluster transcripts are exported via specialized machinery as piRNA precursors (ElMaghraby et al., 2019; Kneuss et al., 2019). Endonucleolytic cleavage, PIWI loading, and 3’ end trimming give rise to 23∼29-nt mature forms of piRNAs (Han et al., 2015; Hayashi et al., 2016; Izumi et al., 2016; Mohn et al., 2015). In addition to the above-mentioned pathway, germline piRNAs are amplified by ping-pong cycle mediated by cytoplasmic PIWI members, Aubergine (Aub) and Ago3 (Lim and Kai, 2007; Sato et al., 2015; Webster et al., 2015). Aub and Ago3 continue primary/trigger piRNA-directed reciprocal cleavages of sense and antisense TE sequences, and secondary/responder piRNA production from 5’ ends of 3’ cleavage fragments. Because target cleavage occurs after 10 nt from 5’ end of trigger piRNAs, trigger and responder pairs hold 10-nt 5’-to-5’ complementarity. Like TE-related transcripts, a couple of endogenous protein-coding transcripts including *stellate (ste), vasa (vas)*, and *pirate (pira)* can produce piRNAs via ping-pong, being targeted by trigger piRNAs derived from *su(ste), AT-Chx*, and *petrel* clusters, respectively (Adashev et al., 2021; Chen et al., 2021a; Kotov et al., 2019). However, aside from these few examples, germ cells generally disfavor the entry of endogenous protein-coding transcripts to piRNA biogenesis (Chung et al., 2021; Mohn et al., 2015). piRNAs display both TE-related and -unrelated sequences, the balance of which can be controlled during development. Similar to Drosophila, murine fetal testes activate ping-pong and accumulate piRNAs containing TE-related repeat sequences (Aravin et al., 2008). However, during neonatal period when spermatocytes and spermatids accumulate, cluster regions having fewer TE sequences are transcribed and processed into so-called pachytene-piRNAs (Chirn et al., 2015; Girard et al., 2006; Li et al., 2013; Özata et al., 2020). Due to the low TE content, many pachytene-piRNAs display unique sequences mapping only once to their origins in genome. Generated diversity rather than individual identities have been considered important for pachytene-piRNA functions. On the other hand, Drosophila major clusters are enriched with TE sequences, and thus do not favorably contribute to piRNA diversity.

Nonetheless, unique sequence piRNAs can be provided by endogenous protein-coding genes (Robine et al., 2009; Saito et al., 2009). A defining characteristic of these genic piRNAs is their production from transcript 3’ untranslated regions (3’UTRs) (Ishizu et al., 2015, 2019; Sun et al., 2021; Takase et al., 2022). As exemplified by *traffic jam* (*tj*) that is one of representative host genes, 3’UTR-piRNAs bind Piwi and exert regulatory effects in gonadal soma (Saito et al., 2009). Overall, unique sequence piRNAs have been poorly characterized in Drosophila germ cells.

Silencing pathways exhibit adaptation to spermatogenesis and regulate testis-unique RNA metabolism. This is well demonstrated with Ago2 interacting with siRNAs and being central in RNA-based immunity against foreign pathogens (Bronkhorst and Van Rij, 2014; Obbard et al., 2006; Tassetto et al., 2017). However, in testes, a unique set of endogenous (endo-)siRNAs are excised from hairpin transcripts by an RNase III enzyme Dicer2 (Dcr2), and incorporated into Ago2 (Lin et al., 2018; Wen et al., 2015). endo-siRNAs direct endogenous mRNA cleavage and support late spermatogenesis (Czech et al., 2008a; Okamura et al., 2008; Wen et al., 2015). We recently reported a co-chaperone associated with Hsp90 machinery, Cyclophilin 40 (Cyp40), is preferentially expressed in male germline and essential for sperm formation (Iki et al., 2020). Ago2 binds testis-unique miRNAs as well as endo-siRNAs, and Cyp40 increases the miRNA occupancy inside RISCs. Other functions of Cyp40 have remained elusive. On the other hand, in piRNA pathway, Ago3 restricts its expression during germline differentiation and is only detectable in spermatogonia (Nagao et al., 2010). Hence, canonical ping-pong between Ago3 and Aub is supposed to be dampened in spermatocytes, though broadly expressed Aub alone can maintain the basal activity (Quénerch’Du et al., 2016). Collectively, these raise a question of whether and how piRNAs and PIWI proteins support Drosophila spermatogenesis particularly in later stage germ cells.

In this study, we reveal hundreds of endogenous protein-coding genes produce unique sequence piRNAs that function in testicular germ cells. These genic piRNAs differ from known species by their origins in coding sequences rather than in 3’UTRs, and by their fates being loaded onto Aub but not Piwi. Furthermore, different from cluster/TE-piRNAs, neither Rhi-dependent cluster transcription nor Ago3-dependent ping-pong amplification is required for their production. Alternatively, our data demonstrate their biogenesis relies on testis-specialized factors including Cyp40 and Ago2. With regard to the latter, siRNAs and miRNAs serve as previously undocumented triggers of secondary piRNA production. These and other results illustrate how germline piRNA diversity is expanded in Drosophila testes for the control of late spermatogenesis.

## Result

### Drosophila testes produce piRNAs from endogenous protein-coding sequences

To deepen the understanding of piRNA pathways underlying spermatogenesis, we first characterized the genomic origins of piRNAs accumulating in testes of *D. melanogaster*. Analyses of deep-sequencing data prepared in house and available in public revealed 23∼29-nt piRNA-like reads accumulate to the exons of endogenous protein-coding genes to different degrees (Figure 1A and S1A). However, total RNAs contain silencing-irrelevant short fragments such as mRNA decay intermediates, leading to false positive piRNA identification. Considering decay fragment accumulation could correlate with mRNA level, we plotted the abundance of 23∼29-nt RNAs relative to mRNAs in individual genes, and excluded those giving lower values from further analyses (Figure 1A, Y-axis value<10). One of such excluded genes, *eEF1alpha1*, indeed displayed a broad fragment distribution ranging from 18 to 29 nt, possibly reflecting its degradome (Figure 1B, dotted line). In contrast, size profile of reads mapping to remained genes showed a clear peak at 24∼26 nt, indicative of piRNAs (Figure 1B, red line). In addition to size, these reads displayed uridine (U) enrichment at 5’ nucleotide, a feature of piRNAs bound to Piwi or Aub (Figure 1C).

**Figure 1.**
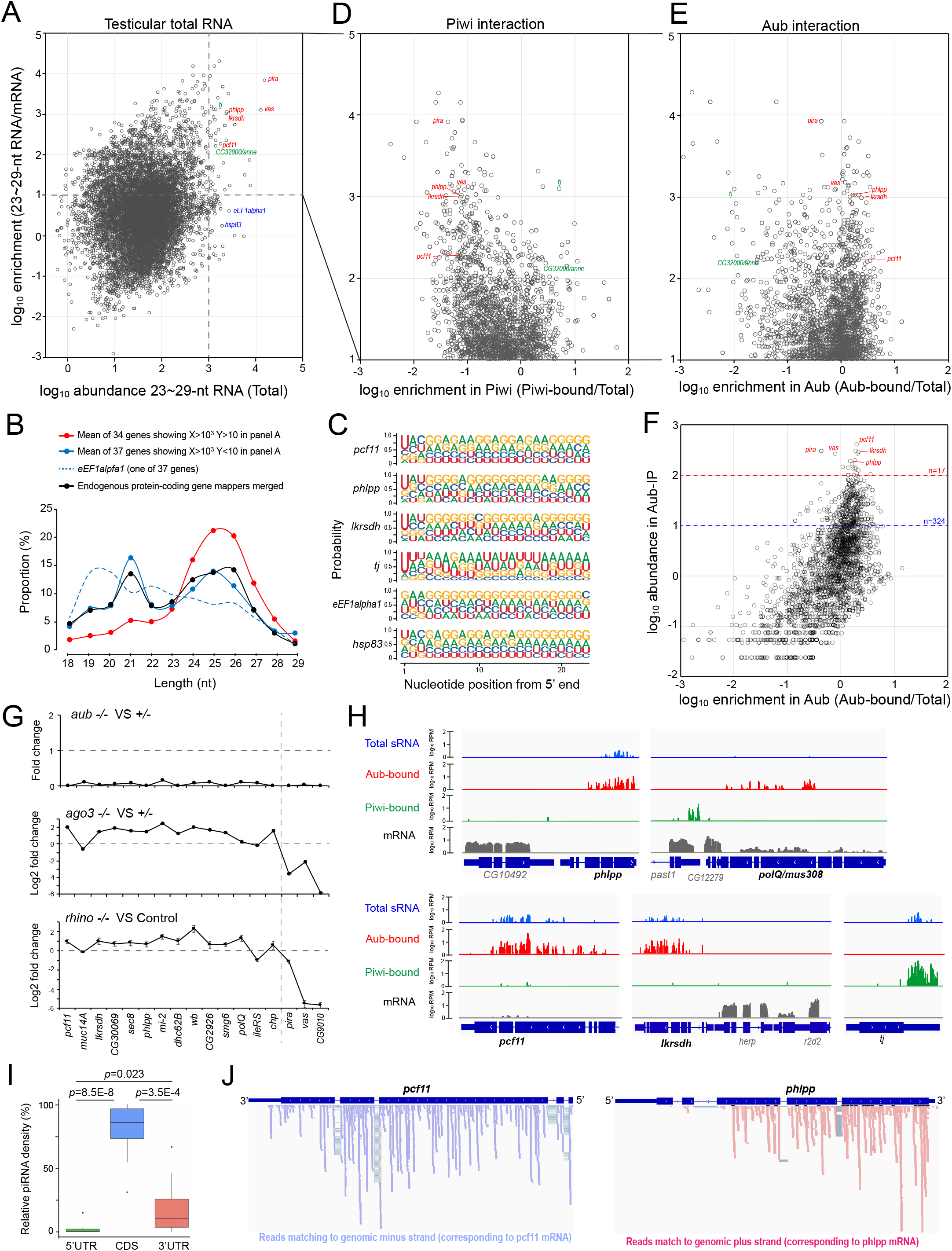
Testicular small RNA mapping to D. melanogaster gene exons. (A) Mapping of 23∼29-nt RNAs accumulating in *D. melanogaster* testes to endogenous protein-coding genes. Abundance (TPM ; transcripts per kilobase million) of 23∼29-nt RNAs mapping to exons (5’UTR + CDS [coding sequence] + 3’UTR) was obtained for individual genes. Mean of 6 data was shown in X-axis. Mean mRNA abundance (TPM) was obtained from 2 data, and enrichment of 23∼29-nt RNAs relative to mRNAs (TPM/TPM) was shown in Y-axis. (B) Size distribution of 18∼29-nt exon-mapping RNAs. (C) Nucleotide probability of 23∼29-nt exon-mapping RNAs. (D, E) Binding of 23∼29-nt RNAs to Piwi or Aub. Abundance (TPM) of 23∼29-nt RNAs in Piwi- or Aub-bound fraction was given as mean of Piwi-IP and GFP-Piwi-IP data, or as mean of Aub-IP and GFP-Aub-IP data. Enrichment (Piwi-bound/total or Aub-bound/total) was shown in X-axis. Genes enriched with 23∼29-nt RNAs relative to mRNAs (TPM/TPM>10, n=2449) were analyzed. (F) Genes producing piRNAs most abundantly for Aub. Abundance (mean RPM [Read per million] of Aub-IP and GFP-Aub-IP) was shown in Y-axis. Seventeen or 324 genes (RPM >100 or >10) were grouped and analyzed later. (G) Loss of *aub, ago3*, or *rhi* effect on exon-mapping piRNAs. Seventeen genes (Aub-IP RPM >100) were analyzed. (H) Origins of piRNAs within gene exons. Bedgraph shows 23∼29-nt RNAs in testis total RNAs (blue), Aub-bound piRNAs (red), Piwi-bound piRNAs (green), and mRNA transcriptome (gray). Gene model shows CDS (bold line) and UTRs (thin line). (I) Density of mapping piRNAs compared between 5’UTR, CDS, and 3’UTR. *pira, vas, CG9010* accumulating *rhi*- and *ago3*-dependent piRNAs were excluded from 17 genes. Remaining 14 genes were analyzed in box plot. *p* (two-tailed paired *t*-test). (J) Strand orientation of Aub-bound piRNAs mapping to *pcf11* and *phlpp*. Arrowhead highlights exon-exon junction reads.

Of those genes enriched with 23∼29-nt RNAs (Figure 1A, Y-axis value>10, n=2449), *traffic jam (tj)* and *CG32000/anne* are known to produce piRNAs interacting with Piwi (Saito et al., 2009). However, others including *phlpp, lkrsdh*, and *pcf11*, have not been linked to piRNA production thus far. Comparison of 23∼29-nt RNA abundance between total and Piwi-bound pools showed that testicular Piwi indeed interacts with piRNAs derived from *tj* and *anne* (Figure 1D). In marked contrast, 23∼29-nt RNAs from many other genes did not exhibit affinity to Piwi, but to Aub alternatively (Figure 1DE). First U bias and 24∼26-nt peak were confirmed for Aub-bound reads mapping to representative genes including *phlpp, lkrsdh*, and *pcf11* (Figure 1F RPM>100 n=17 and S1BC). Moreover, these piRNA-like products were nearly depleted in total RNAs of *aub* mutant testes (Figure 1G). Taken together, these data indicate endogenous protein-coding gene exons produce *bona fide* piRNAs interacting with Aub in testicular germ cells.

Of 17 genes producing piRNAs most abundantly, *vas* and *pirate (pira)* are known to be targeted by piRNAs from *AT-ChX* and *petrel* clusters, respectively; secondary piRNAs can be generated from their cleaved mRNAs, like TEs and cluster transcripts (Chen et al., 2021a; Kotov et al., 2019). Consistently, the levels of *vas-* and *pira*-derived piRNAs showed severe decrease in testes lacking *rhi* or *ago3*, that is crucial for cluster transcription or ping-pong amplification, respectively (Figure 1G). In sharp contrast, piRNAs from other 14 genes excluding *CG9010* did not decrease their abundance, implying a distinct mechanism underlying their biogenesis. We further found piRNAs from these 14 members were predominantly mapped to protein-coding regions, a feature differed from known genic species mainly derived from 3’UTRs (Figure 1HI). Considering these characteristics, we hereafter refer these piRNAs, representatively produced by aforementioned 14 genes, as <u>endo</u>genous protein-coding sequence-derived (endo-)piRNAs. endo-piRNAs showed a near uniform sense-strand orientation corresponding to mRNAs, with an exception of *muc14A* dominated with antisense reads (Figure S1DE). In addition, endo-piRNAs were rarely derived from introns, and a fraction of reads contained exon-exon junction sequences (Figure 1J). These features suggest that spliced, sense-strand transcripts including mRNAs can serve as the precursors.

We examined if endo-piRNA production is regulated during germline development. Re-analysis of small RNAs present in testes lacking germline differentiation factors (Figure S1F) (Quénerch’Du et al., 2016) showed endo-piRNAs are barely detectable in early spermatogonial cells, but can be seen in primary spermatocytes (Figure S1F). These results suggest that endo-piRNA production is induced as germ cells differentiate from spermatogonia to spermatocytes, a pattern opposed to cluster/TE-piRNA production.

### endo-piRNAs are preferentially produced in testes compared to ovaries

To examine if ovaries can produce endo-piRNAs like testes, we mined the publicly available data of total small RNAs and Aub-bound piRNAs in ovaries, and compared with testicular data sets (Figure S2A). Curiously, ovarian Aub did not accumulate endo-piRNAs mapping to genes identified in testes (Figure 2A). Moreover, *de novo* screening of genes hosting endo-piRNAs using ovarian data sets listed up much fewer candidates compared to those in testes (Figure S2BC). On the other hand, by genome-wide profiling, we did not observe significant difference in the proportion of exon mappers between testicular and ovarian data, possibly due to non-piRNA fragment contamination (Figure S2D). Overall, these results suggest endo-piRNAs are preferentially produced in testes rather than ovaries.

**Figure 2.**
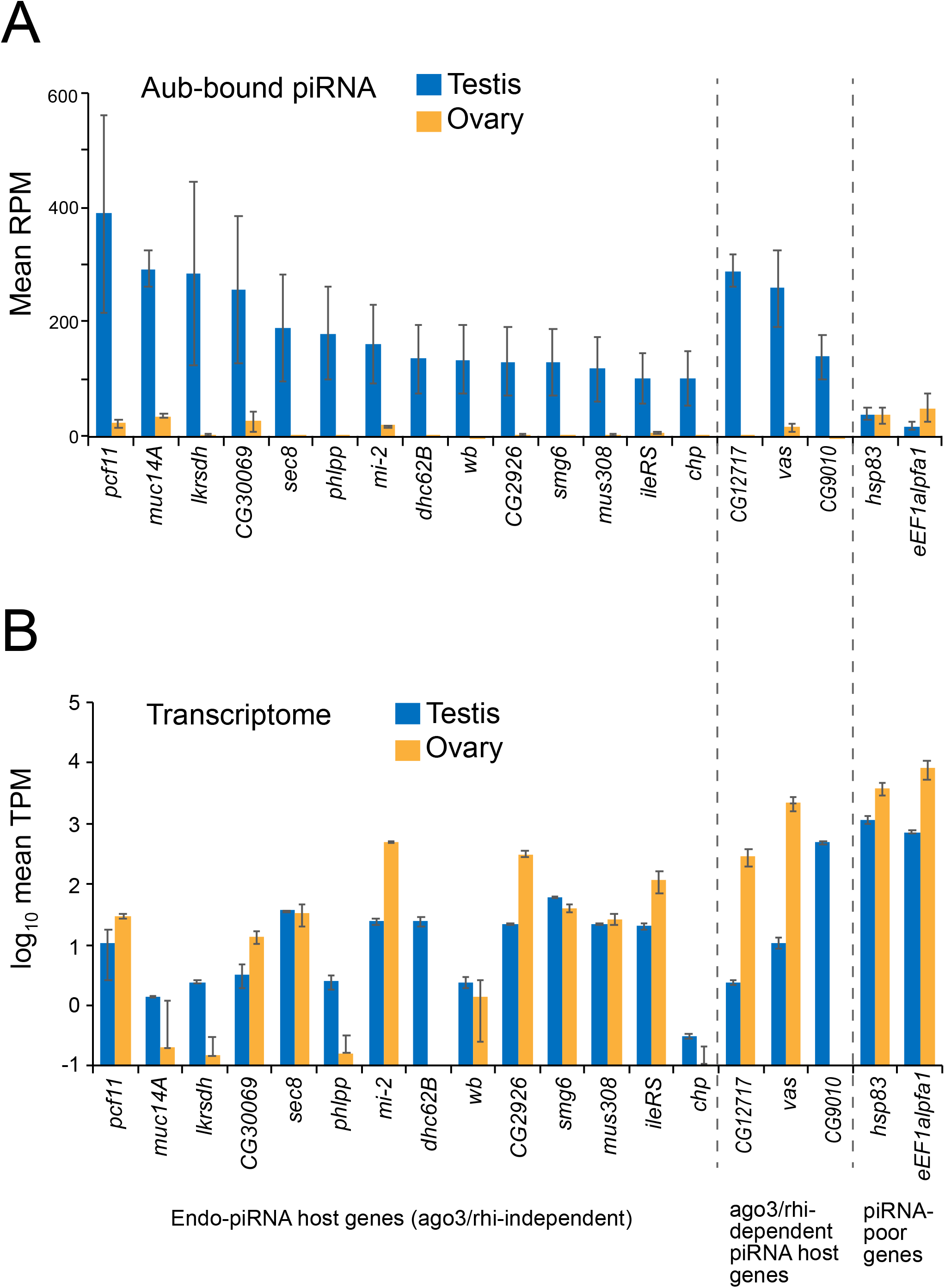
endo-piRNAs and cognate mRNAs present in testes and ovaries. (A) endo-piRNA abundance (RPM) in Aub-RISCs or (B) cognate transcript abundance (TPM) compared between testes and ovaries. Indicated genes include 17 representative piRNA producers in testes, together with 2 piRNA-poor references (*hsp83* and *eEF1alpha1*). Mean ± standard deviation (s.d.) of 2 data set was shown.

Contrary to endo-piRNAs, the expression levels of many, if not all, host gene mRNAs were comparable between ovaries and testes (Figure 2B). Therefore, lessor accumulation of Aub-bound endo-piRNAs in ovaries would not be solely attributable to the fewer amount of precursor transcripts. Rather, these results imply testicular germ cells express factors that can act on the selected protein-coding transcripts and enable their entry to piRNA biogenesis pathway.

### Aub is physically linked to Cyp40, an Hsp90 cochaperone specialized for spermatogenesis

What testicular factors can activate endo-piRNA production and Aub loading? Small RNA loading step in RISC formation relies on Hsp70/Hsp90 chaperone machineries (Iki et al., 2010; Iwasaki et al., 2010; Izumi et al., 2013; Miyoshi et al., 2010). In Drosophila, an Hsp90 co-chaperone Cyp40 is expressed in testicular germ cells and essential for sperm formation, however, its substrates/clients are elusive besides Ago2 (Iki et al., 2020). Hence, we performed Cyp40 client screening based on proximity-dependent biotin identification (BioID/TurboID) (Branon et al., 2018; Roux et al., 2012), that could be advanced in identifying transient chaperone-client interactions compared to conventional immunoprecipitation analyses (Figure 3A). Cyp40 fused to an engineered biotin ligase termed mini (m)Turbo was expressed in germ cells using UASp-Gal4 system (Figure 3A) (Rørth, 1998). mTurbo-Cyp40 rescued the defective spermatid differentiation and sperm storage in *cyp40*-null testes, confirming the functionality of fusion construct (Figure S3A). In denaturing gel electrophoresis, potentially biotinylated proteins were detected as distinct bands when Cyp40, but not GFP or nonfunctional Cyp40 variant (RKAA or *Δ*TPR) was used as bait (Figure 3B) (Iki et al., 2020).

**Figure 3.**
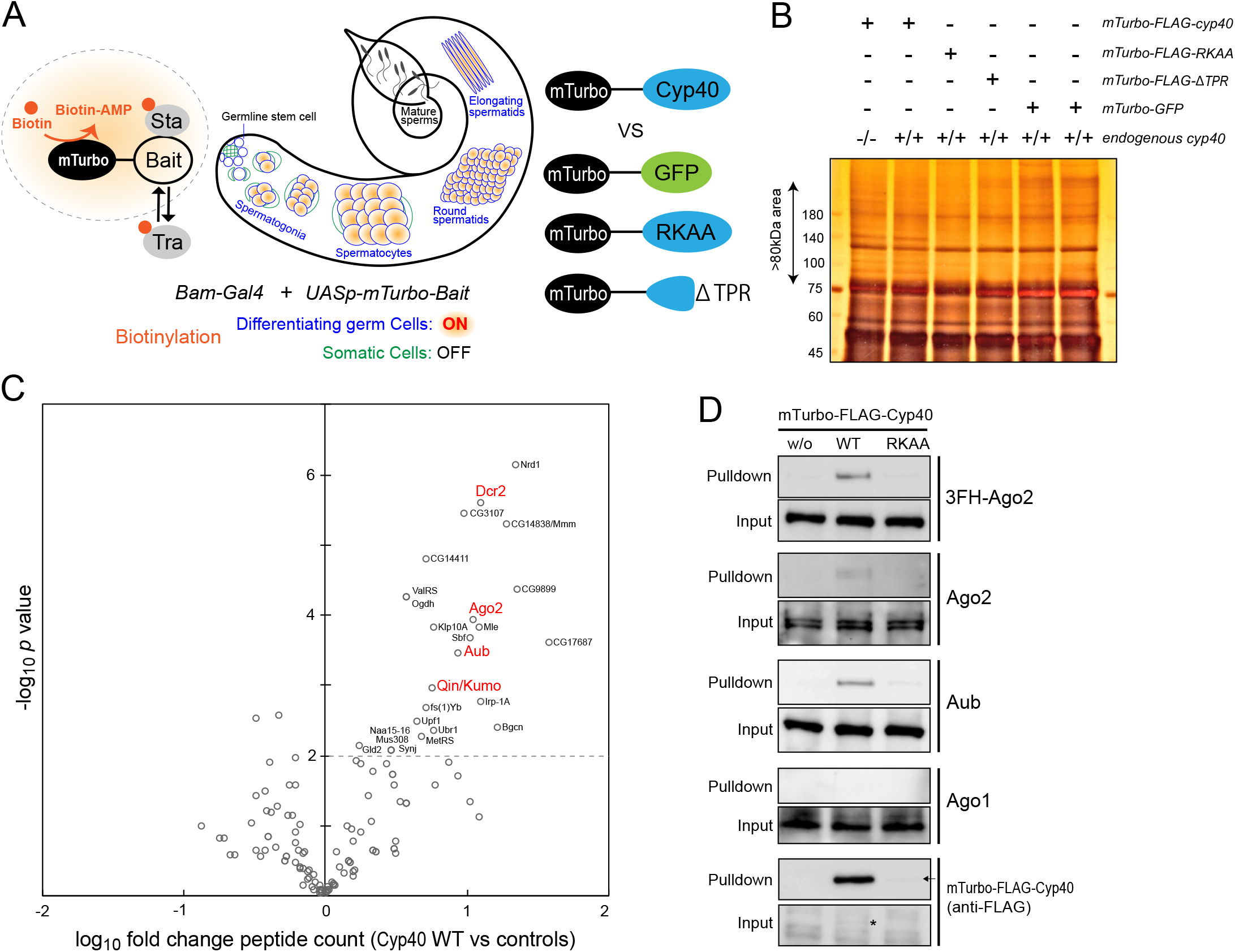
Cyp40 client screening in testicular germ cells using TurboID. (A) Principles of germline BioID/TurboID developed in this study. Using free biotins, an engineered biotinylation enzyme called mini(m)Turbo generates diffuse pattern of biotin-AMP, which adds biotins to lysine residues of proteins close to the selected bait. Not only stable (Sta) but also transient (Tra) interactors such as chaperone clients can be biotin-labeled. mTurbo was fused to bait proteins including Cyp40, non-functional Cyp40 derivatives (RKAA or *Δ*TPR), or GFP. RKAA and *Δ*TPR lack affinity to Hsp90. (B) Silver staining of proteins purified with streptavidin. Cyp40-dependent signals were enriched in >∼80kDa area from which peptides were extracted and analyzed by mass spectrometry. (C) Volcano plot summarizing mass-spectrometry data. Spectrum counts of proteins purified from testes expressing mTurbo-FLAG-Cyp40 (2 data; in the presence or absence of endogenous *cyp40*) were compared with those in control conditions (4 data; single replicate of mTurbo-FLAG-Cyp40^RKAA^ and mTurbo-FLAG-Cyp40^*Δ*TPR^, duplicate of mTurbo-GFP). X-axis; enrichment of spectrum counts (Cyp40/control), Y-axis; *p* (two-tailed unpaired *t*-test). Names of significantly (*p*<0.01) enriched proteins were indicated. (D) Immunoblotting of Ago proteins and mTurbo-FLAG-Cyp40 present in testes (Input) and purified with streptavidin (Pulldown). For Ago2, both 3FLAG-HA-tagged (3FH-)Ago2 expressed under native promoter activity and endogenous Ago2 were analyzed. Of note, mTurbo-FLAG-Cyp40 proteins were barely detectable in input (indicated by asterisk), but strongly enriched by streptavidin pulldown, indicating efficient self biotinylation. Steady state level of mTurbo-FLAG-Cyp40^RKAA^ was much lower and only visible in pulldown fraction (Arrow).

The screening identified dozens of factors in the physical proximity of Cyp40. Consistent with our previous findings, these included Ago2 and, in addition, its essential partner Dcr2 (Figure 3C and S3B) (Iki et al., 2020). Intriguingly, Aub was identified as one of proximity factors and candidate clients of Cyp40. Moreover, our profiling listed up Qin/Kumo, a Tudor domain protein associated with Aub-RISC formation (Anand and Kai, 2012; Zhang et al., 2011). Contrastively, no significant signal enrichment was seen for other Ago members. Immunoblotting using respective Ago antibodies confirmed the mass spectrometry data (Figure 3D and S3C). Of note, Cyp40-based TurboID did not list up Hsp90, the partner of Cyp40 in chaperone machinery and thus an expected biotinylation target. This perhaps implicates the relative distance between mTurbo and Hsp90 in complexes or other restrictions of employed strategy. Nevertheless, the unbiased screening revealed the selective link of Cyp40 to Aub and Ago2 among Ago family members.

### Aub loading with endo-piRNAs is selectively assisted by Cyp40

Above results prompted us to explore a possibility that Cyp40 is involved in Aub-dependent piRNA pathways in testes. Given chaperone functions are associated with RISC formation, Cyp40 could affect piRNAs inside Aub-RISCs. To examine this possibility, Aub-bound piRNAs in *cyp40* null testes were analyzed by deep-sequencing and compared with those of heterozygous siblings. Aub protein level was comparable between those samples (Figure 4A and S4A) and, accordingly, the abundance of piRNAs mapping to cluster or TE sequences were largely unaffected (Figure 4B and S4B). In sharp contrast to cluster/TE-piRNAs, however, exon mappers enriched with endo-piRNAs (exons of 324 genes, Figure 1F) were reduced in *cyp40* mutant testes (Figure 4B, fold change [FC] = 0.71±0.05). Individual analyses of 324 genes confirmed the general trend of mapping-reads reduction (Figure 4C, Table S1, FC = 0.73±0.18). Of those, *pira* was one of exceptions; like cluster/TE-piRNAs, its piRNA level was unaffected (Figure 4D). In sum, *bona fide* endo-piRNAs derived from 14 genes (Figure 1G) exhibited a severe decrease in abundance upon loss of *cyp40* from testes (Figure 4D, FC = 0.59±0.12). These results suggest that endo-piRNA loading to Aub is selectively assisted by Cyp40. The uncovered *cyp40*-dependency of endo-piRNA accumulation, together with the *rhi/ago3*-independency (Figure 1G), supports the idea that endo-piRNAs and cluster/TE-piRNAs are produced via distinct mechanisms.

**Figure 4.**
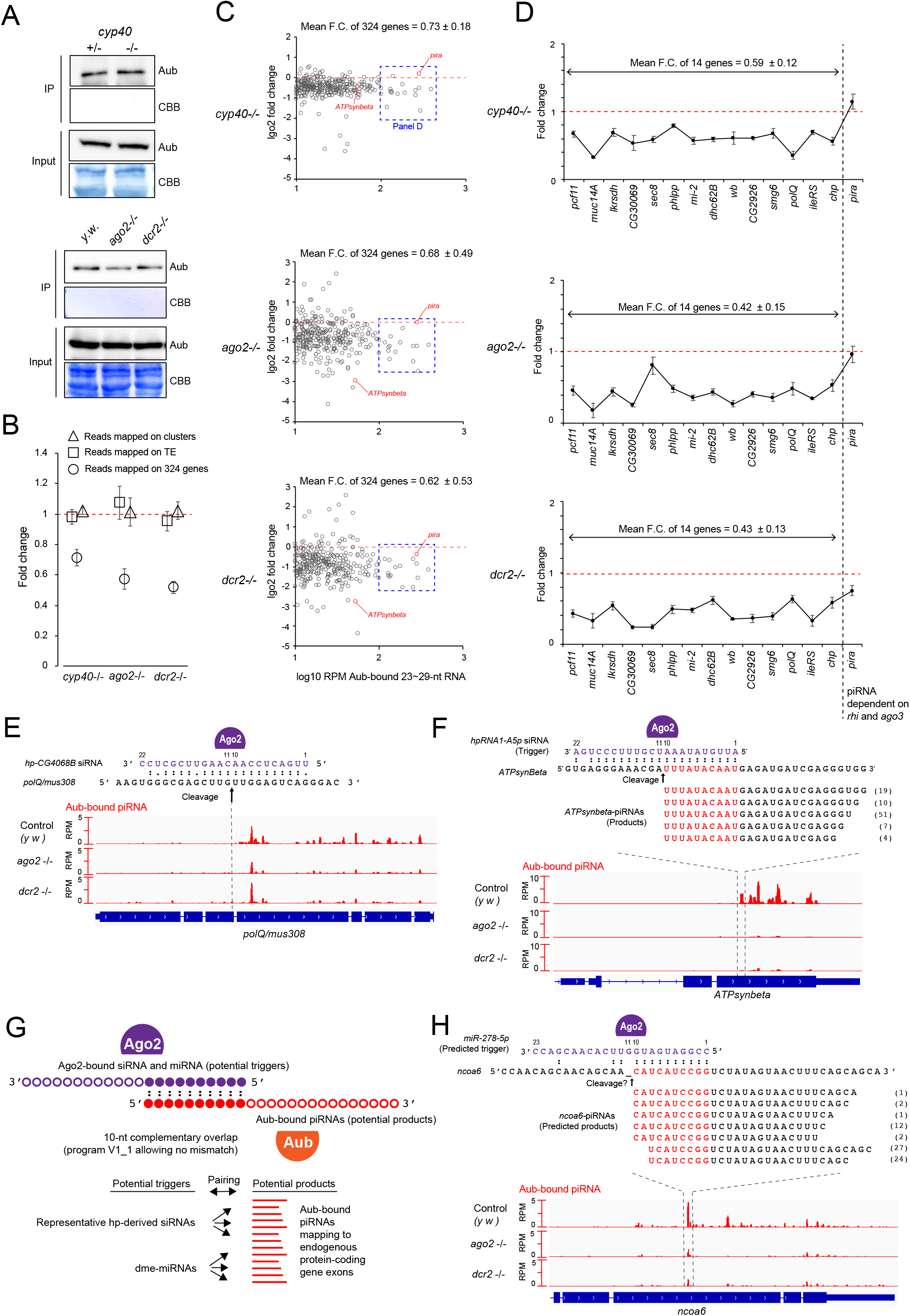
Molecular requirement for endo-piRNA biogenesis. (A) Immunoblotting of endogenous Aub proteins present in testes (Input) and immunopurified with anti-Aub antibodies for deep-sequencing (IP). *cyp40*^*KO/Df*^ null mutant (-/-) was compared to the sibling *cyp40*^*KO*/*TM6*^ heterozygous control (+/-). *ago2*^*454/Df*^ and *dcr2*^*L811fsX/811fsX*^ null mutants (-/-) were compared to *y w* control testes. Coomassie brilliant blue (CBB) staining serves as protein loading control. (B) Effect of *cyp40, ago2*, or *dcr2* loss on piRNA abundance (RPM) inside Aub-RISCs. Aub-bound piRNAs were grouped as cluster mappers, TE mappers, or exon mappers. For exon mappers, selected 324 genes were analyzed to enrich endo-piRNAs (Aub-IP RPM>10, see Figure 1F). Fold changes were given by *cyp40* mutant/heterozygous, *ago2* mutant/*y w*, and *dcr2* mutant/*y w* comparisons, respectively. Mean ± s.d. of 2 data set was shown. (C, D) Effect of *cyp40, ago2*, or *dcr2* loss on exon mappers in individual genes. piRNA-enriched 324 genes (C), and 14 (+1=*pira*) genes (D) were analyzed. *pira* serves as a reference producing piRNAs in a *ago3*/*rhi*-dependent manner, unlike endo-piRNAs. (E, F) siRNA target sites and piRNA origins on *mus308/polQ* and *ATPsynbeta*. In *ATPsynbeta*, targeting siRNAs and produced piRNAs exhibit 5’-to-5’ 10-nt complementary overlap. Numbers next to sequences indicate read counts in merged Aub-IP data (*y w*). (G) Screening of 5’-to-5’ 10-nt complementary overlap. piRNAs paired with siRNAs or miRNAs were searched within Aub-bound exon mappers. (H) Ago2-sorted *miR-278-5p* and *ncoa6*-derived piRNAs predicted as trigger-product pairs.

### Ago2 and binding small RNAs activate endo-piRNA production

endo-piRNAs often exhibit mapping preference within the protein-coding regions, as being exemplified by *DNA polymerase theta (polQ)/mus308* and *ATP synthase beta* (*ATPsynbeta)*. Transcripts of these genes are known targets of siRNAs derived from *CG4068* and *hpRNA1*, respectively, and cleaved by Ago2 (Figure 4EF) (Czech et al., 2008b; Okamura and Lai, 2008; Wen et al., 2015). For both, piRNAs were near exclusively mapped downstream of siRNA target sites. Particularly for *ATPsynbeta*, a fraction of piRNAs were produced precisely from the 5’ end exposed by cleavage, and thus targeting siRNAs and produced piRNAs hold 5’-to-5’ complementary overlap over 10 nt (Figure 4F).

These observations imply that Ago2 activities may trigger endo-piRNA production. To examine this, Aub-bound piRNAs in *ago2* or *dcr2* null mutant testes were analyzed and compared with those of the control. Similar to *cyp40* mutant analyses, loss of *ago2* or *dcr2* did not alter the abundance of Aub proteins and cluster/TE-piRNAs (Figure 4AB and S4B), but results in severe reduction of endo-piRNA-enriched exon mappers (Figure 4B, FC = 0.57±0.06 and 0.53±0.04). Individual genes exhibited more variable effects (Figure 4C, FC = 0.68±0.49 and 0.62±0.53), but nonetheless there was correlation between *ago2*- and *dcr2*-deficient conditions (Figure S4C, *r* = 0.48). To this end, genuine endo-piRNAs from 14 genes showed most severe decrease (Figure 4D, FC = 0.42±0.15 and 0.43±0.13).

Our data highlight that endo-piRNAs from a set of genes including *ATPsynbeta* were nearly depleted in testes lacking Ago2-Dcr2 pathway, however, these extreme cases were rare (Figure 4CF and S4C). In addition, our 10-nt overlap screening did not find an siRNA pairing with endo-piRNAs other than *hpRNA1*-siRNA, but predicted potential trigger functions of Ago2-bound miRNAs (Figure 4GH, S4D, Table S2). Therefore, we conclude that Ago2 activities relying on siRNAs and miRNAs are in general important, if not essential, for endo-piRNA production.

### *aub* is crucial for spermatid differentiation and male fertility

As a step toward understanding the biological roles of endo-piRNA pathway, we examined if and how *aub* is required for spermatogenesis. A loss-of-function allele, *aub*^*N11/HN2*^, contained much fewer sperms in seminal vesicles compared to the heterozygous siblings, and the males were nearly sterile (Figure 5AB and S5A). Close examination of spermatogenesis revealed that although spermatogonia and spermatocytes were largely unaffected (Figure S5B), individualization complexes (ICs) formed by spermatids were severely disorganized (Figure 5AC). Those defects were largely recovered by introduced *aub* transgene, indicating that the loss of *aub* is causal for the phenotypes. Like *aub, ago2* is important for endo-piRNA production (Figure 4), and plays key roles in late spermatogenesis (Figure S5BC) (Wen et al., 2015). In addition, double knock-out (dKO) of *aub* and *ago2* did not exhibit any discernible defects in spermatogonia and spermatocytes, like single KOs (Figure S5BC). These results suggest endo-piRNA pathway involving both Aub and Ago2 can support late spermatogenesis.

**Figure 5.**
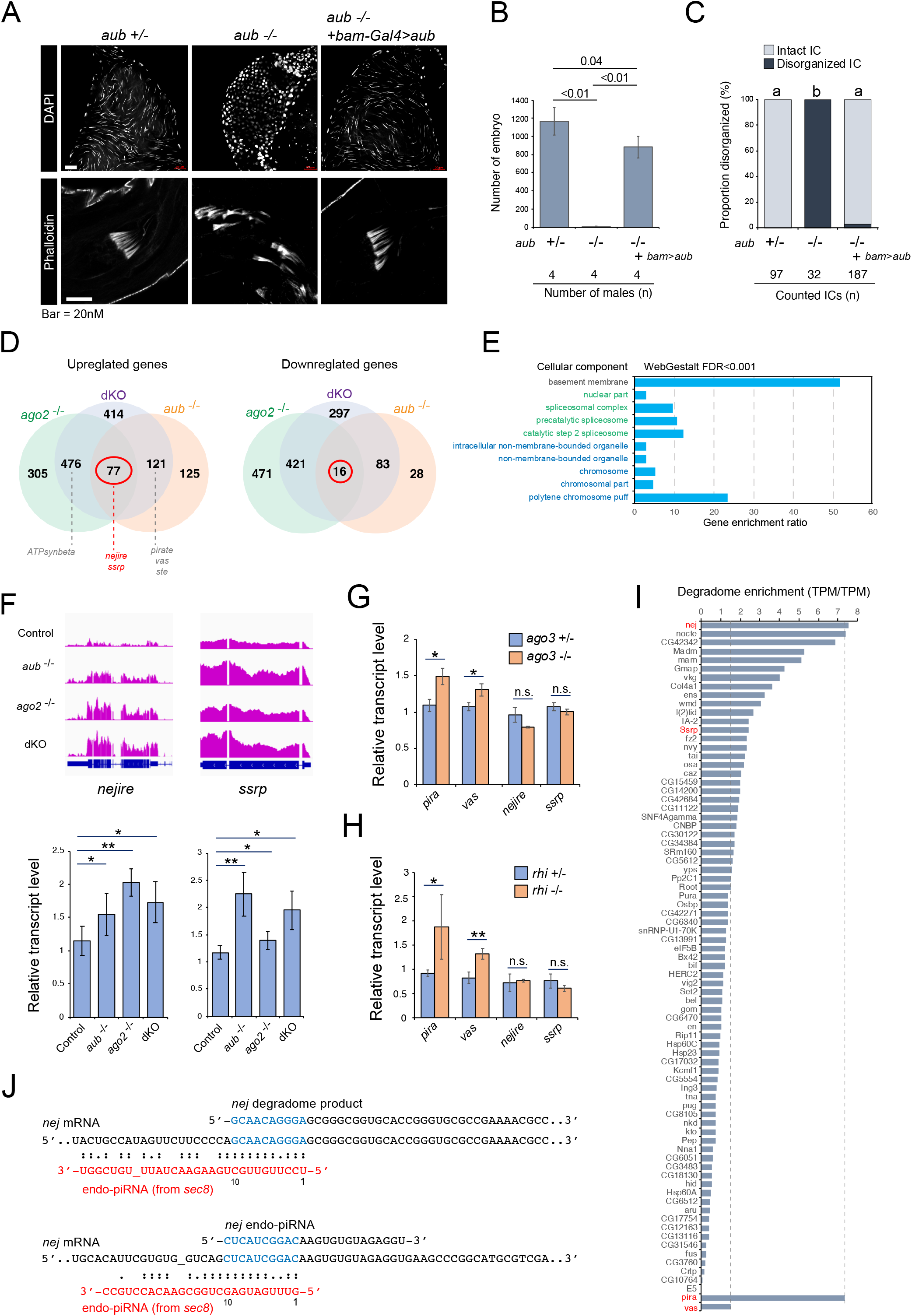
Functions of Aub, Ago2, and endo-piRNAs in late spermatogenesis. (A) Defective spermatogenesis in testes lacking *aub* (-/-; *aub*^*N11/HN2*^), and the rescue by *aub* transgene (*bam-Gal4>UASp-aub*). *aub* heterozygous siblings (+/-) serve as control. Sperm nuclei stored in seminal vesicles (DAPI). Individualization complexes (ICs) in elongating spermatids (Phalloidin). (B) Male fertility test. Mean ± s.d. was shown. *p* (two-tailed unpaired t-test) were indicated. (C) Disorganized IC counting. Differential characters indicate statistically significant difference (*p*<0.01) in Tukey’s test. (D) Venn diagram highlights genes regulated by both *aub* and *ago2*. Transcriptome of *aub*^*N11/HN2*^ mutant (*aub*^*-/-*^ *ago2*^*+/-*^), *ago2*^*454/Df*^ mutant (*aub*^*+/-*^ *ago2*^*-/-*^), or double knock-out (dKO) (*aub*^*-/-*^ *ago2*^*-/*-^) was compared to that of sibling control (*aub*^*+/-*^ *ago2*^*+/-*^). Analysis was based on biological duplicates. Mis-regulated genes were extracted by FDR<0.0001. (E) Gene ontology (cellular components) analyses of overexpressed 77 genes. (F) Deep-sequencing profile (bedgraph) and qPCR measurement (bar graph) of *nej* and *ssrp* transcripts. Mean ± s.d. was shown. Asterisk(s); single (*p*<0.05) double (*p*<0.01) in two-tailed t-test. (G, H) qPCR measurement of gene transcripts in *ago3*^*t2/t3*^ (-/-) or *rhi*^*02086/Df*^ (-/-), compared to respective heterozygous sibling controls (+/-). (I) Enrichment of degradome products relative to polyadenylated transcripts (TPM/TPM). Genes overexpressed in *aub* and *ago2* mutants (n=77) were analyzed. *pira* and *vas* serve as reference target of piRNAs for cleavage. (J) Signatures of endo-piRNA-directed *nej* mRNA cleavage. 5’-to-5’ 10-nt complementary overlaps between targeting endo-piRNAs and degradome products (top), and between targeting endo-piRNAs and produced endo-piRNAs (bottom).

### endo-piRNAs can exert regulatory effect on endogenous gene expression

endo-piRNAs are enriched with TE-unrelated sequences, and their impacts on TEs are suspicious. In addition, since their sense-strand sequences match to host mRNAs, *cis* regulatory effects are also unlikely. Considering these, and to examine the gene-regulatory functions of endo-piRNAs, we profiled the transcriptome of testes deficient for endo-piRNA production. Testes lacking *aub (aub*^*-/-*^ *ago2*^*+/-*^*), ago2 (aub*^*+/-*^ *ago2*^*-/-*^), or both (dKO; *aub*^*-/-*^ *ago2*^*-/-*^) were obtained from progenies derived from the same parents, together with those of sibling heterozygous controls. Genes controlled by endo-piRNA pathway could be mis-regulated in all mutant conditions. Consistent with previous studies, *ATPsynbeta*, a direct target of *hpRNA1*-derived siRNAs, was identified as one of overexpressed genes in *ago2* mutant and dKO testes, but not in *aub* mutants (Figure 5D and S5D). In addition, cluster-piRNA targets including *ste, pira*, and *vas*, were derepressed in *aub* mutant and dKO testes, but not in *ago2* mutants. We then listed up genes mis-regulated in both *aub* and *ago2* mutants. The list contained 77 upregulated and 16 downregulated genes (Figure 5D, FDR<0.0001), and all were mis-regulated in the same way in dKO. Gene ontology analyses indicated that overexpressed genes are significantly enriched with those functions associated with chromosome, splicing, and basement membrane (Figure 5E, FDR<0.001). Quantitative PCR (qPCR) confirmed the transcriptome output by analyzing *nejire (nej)/CBP* and *structure-specific recognition protein* (*ssrp)*, whose products participate in chromatin regulation (Figure 5F). Of note, *nej* encoding histone acetyltransferase is involved in spermatid differentiation and male fertility (Hundertmark et al., 2018). Loss of *rhi* or *ago3* did not affect *nej* and *ssrp* levels, thus their regulation by cluster/TE-piRNA pathway is unlikely (Figure 5GH). endo-piRNAs may recognize and direct the cleave of *nej* and *ssrp* transcripts (Figure S5E). Indeed, degradome profiling revealed both genes are enriched with 5’ monophosphate fragments that can be made by endonucleolytic cleavage (Figure 5I). An extreme case is *nej*, whose degradome enrichment was comparable to that of *pira* tightly regulated by cluster-piRNAs via cleavage (Chen et al., 2021a). To support the idea that *nej* cleavage is directed by endo-piRNAs, 5’-to-5’ 10-nt complementary overlap was identified between targeting endo-piRNAs and degradome products (Figure 5J). Notably, *nej* itself is one of 324 genes producing piRNAs, and we also found 5’-to-5’ 10-nt overlap between targeting endo-piRNAs and produced endo-piRNAs, implying their cleavage trigger-responder relationship (Figure 5J). Taken together, these results suggest that endo-piRNAs regulate endogenous genes in late spermatogenesis, at least in part via direct mRNA cleavage.

### Mis-regulated histone H4 acetylations and failure of histone-to-protamine transition in the absence of Aub and interacting piRNAs

In many animals, differentiating spermatids remodel their chromatins by replacing most histones with smaller basic proteins called protamines (Balhorn, 2007). Histone modifications including H4 hyperacetylation are associated with histone-to-protamine transition (Awe and Renkawitz-Pohl, 2010; Dottermusch-Heidel et al., 2014; Goudarzi et al., 2014). Given that *nej* is regulated by endo-piRNA pathway (Figure 5), histone acetylation and associated transition events could be impaired by loss of *aub*. Consistent with previous studies (Hundertmark et al., 2018), acetylated lysine (K) signals on H4 were clearly detectable in spermatid nuclei (Figure S6 and 6AB). During transition, reducing signals of H4 and the acetylated K5, K8, and K12 could overlap with accumulating protamine B (protB), but it lasts only transiently (Figure 6B, upper panels). Later, ProtB-positive and H4-negative nuclei sharpened their structures and started individualization. We found *aub* mutant testes maintained higher levels of H4K8Ac and H4K12Ac, known *nej*-sensitive modifications (Ludlam et al., 2002), compared to heterozygous control and transgene rescue conditions (Figure 6C). In addition, *aub* mutant testes accumulated individualized nuclei that abnormally contain both ProtB and acetylated H4 (Figure 6B, lower panels). These results are in line with the notion that regulation of *nej* and other genes by endo-piRNA pathway underlies proper chromatin remodeling and individualization during spermatid differentiation.

**Figure 6.**
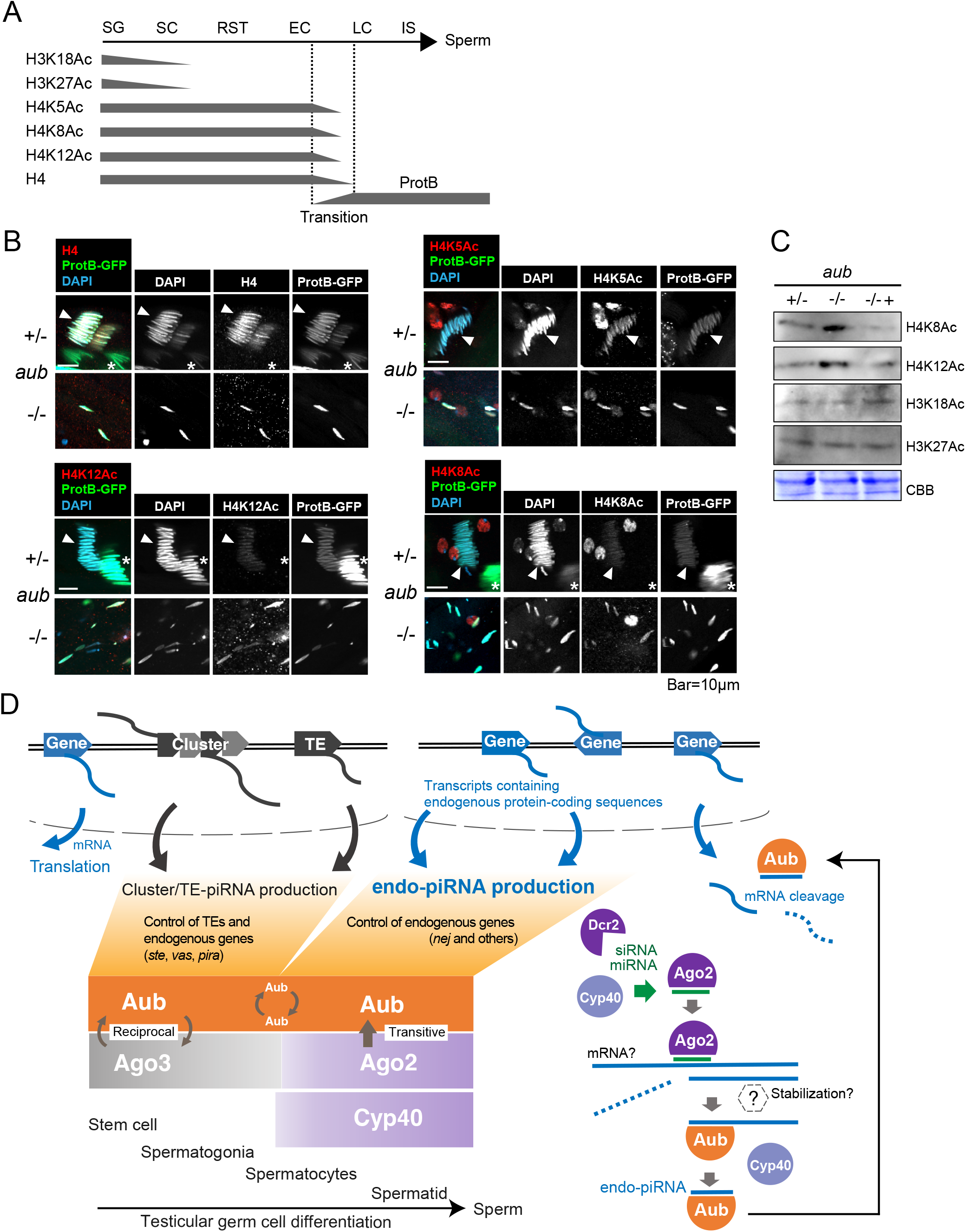
A role of Aub in histone-to-protamine transition, and a model of piRNA biogenesis in testicular germ cells. (A) Summarized histone H3 and H4 acetylation patterns during spermatogenesis. SG; spermatogonia, SC; spermatocytes, RST; round spermatids, EC; early canoe stage spermatids, LC; late canoe stage spermatids, IC; individualizing spermatids. (B) Spermatid nuclei in testes of *aub* heterozygous control (+/-) or *aub*^*N11/HN2*^ null mutants (-/-). Arrowhead; nuclei under transition, both H4 and ProtB signals co-exist. Asterisk; nuclei after transition and before individualization, H4 and associated acetylations are undetectable. (C) Immunoblotting of acetyl histone H3 and H4 extracted from testes of *aub* heterozygous control (+/-) or *aub*^*N11/HN2*^ null mutants (-/-) in the absence or presence of *aub* transgene. CBB serves as protein loading control. (D) Model of germline piRNA biogenesis and functions in Drosophila testes. In spermatogonia and early spermatocytes, many piRNAs are produced from cluster transcripts and TE mRNAs via ping-pong amplification. From primary spermatocyte stages when Ago3 expression and ping-pong activity are reduced, Aub is able to be loaded with fragments containing endogenous protein-coding sequences, forming RISCs containing endo-piRNAs. Cyo40 and Ago2 activities are required for normal endo-piRNA accumulation inside Aub-RISCs. Most endo-piRNAs display unique sequences and thus expand germline piRNA diversity in testes. endo-piRNA-directed mRNA cleavage can support late spermatogenesis.

## Discussion

Our bioinformatic, proteomic, and genetic approaches revealed a unique pathway in which hundreds of selected endogenous protein-coding genes contribute to non-coding piRNA biogenesis by providing precursors to Aub. Namely, endo-piRNA production involves testis-specialized factors including Cyp40 and Ago2, but not Ago3 and Rhi, and hence, the pathway is distinguished from that of cluster/TE-piRNAs. How can two distinct piRNA biogenesis pathways cooperate in testicular germ cells? In testes, Ago3 is restricted in spermatogonia (Nagao et al., 2010; Quénerch’Du et al., 2016), and Rhi loses its expression in early spermatocyte differentiation (Chen et al., 2021b). Hence, cluster/TE-piRNA biogenesis is supposed to be weakened as germ cells differentiate into spermatocytes. On the other hand, Aub and Cyp40 are broadly expressed (Iki et al., 2020; Quénerch’Du et al., 2016), and our data suggest endo-piRNA production is induced in spermatogonia-to-spermatocytes transition. Taken together, we propose that spatiotemporal segregation of two mechanisms generate the full repertoire of germline piRNAs during Drosophila spermatogenesis (Figure 6D). Thus far, only piRNAs have been characterized as triggers of secondary piRNA biogenesis (Ozata et al., 2019). This study expanded the trigger list by adding siRNAs and perhaps miRNAs (Figure 4). However, the conservation of non-piRNA-directed secondary piRNA biogenesis across animals is currently uncertain. In many organisms including plants, fungi, and different animals, transcripts targeted by trigger small RNAs recruit RNA-dependent RNA polymerase (RdRp) enzymes, whose activities mediate silencing amplification through secondary siRNA biogenesis. In plants, mi/siRNA-directed transcript cleavage can recruit RdRp, and produced dsRNAs are processed by RNase III enzymes into secondary siRNAs (Liu et al., 2020). In nematodes, transcripts targeted by primary piRNAs recruits RdRp. Synthesized short complementary fragments are directly incorporated into WAGO subfamily members as secondary, so-called 22G-siRNAs (Weiser and Kim, 2019). *RdRp* genes can be found in the genomes of animals not restricted to nematodes. However, loss of *RdRp* has occurred frequently and independently during evolution, resulting in the sporadic absence of RdRp activities in organisms including Drosophila and mammals (Pinzón et al., 2019). In insects, house dust mites have lost their piRNA pathway factors, which might reflect the functional replacement by RdRp-dependent siRNA modules (Mondal et al., 2018). The prevalence of si/miRNA-directed piRNA biogenesis, an RdRp-independent mechanism to amplify and diversify small RNAs, might have relevance to the absence of *RdRp* in genome.

An important aspect in secondary/responder small RNA biogenesis is to protect the precursor transcripts from degradation and to promote their entry to processing pathways. In ping-pong amplification, a Tudor-domain protein Krimper tethers piRNA-bound Aub and piRNA-free Ago3 in physical proximity to facilitate the efficient transfer of cleaved transcripts to Ago3 (Sato et al., 2015; Webster et al., 2015). In phased piRNA biogenesis, cleaved fragments are carried by RNA helicase Armitage to the processing center on mitochondrial outer membrane (Ge et al., 2019; Ishizu et al., 2019). On the other hand, in plant cells where secondary siRNAs accumulate, 3’ cleavage fragments and RdRp templates are stabilized by RISCs and associated factors (Sakurai et al., 2021; Yoshikawa et al., 2021). The above evidence lets us to hypothesize Drosophila testicular germ cells may maintain mechanism(s) promoting the transfer of precursors to Aub for efficient endo-piRNA production. Responsible factors might exist among those identified in the physical proximity of Cyp40 (Figure 3), or could be identified in the interactome of Aub or Ago2 in future studies. It has been shown that, in plants, precursors can be stabilized by selected small RNAs, which are distinguished by their length of 22 nt but not the shorter isoforms (Chen et al., 2010; Cuperus et al., 2010; Iki et al., 2018; Yoshikawa et al., 2013, 2021). We mention the major form of *CG4068* and *hpRNA1* siRNAs associated with endo-piRNA production is exceptionally 22 nt in length (Figure 4), which are distinct from the most abundant 21-nt ones in testicular pools. Perhaps not all siRNAs are capable of inducing endo-piRNA production. In addition, our data do not exclude the possible involvement of Ago1, interacting with vast majority of miRNAs and a fraction of siRNAs, in endo-piRNA biogenesis. It is an open question whether any Ago- or small RNA-selective rule can be applied to define endo-piRNA triggers.

Why do testes produce endo-piRNAs? Though we provided evidence that *nej* is one of the direct targets of endo-piRNAs (Figure 5), the global view of endo-piRNA mode of actions during late spermatogenesis, when Aub exerts crucial functions, awaits further investigation. Murine pachytene-piRNAs are expressed in neonatal testes accumulating spermatocytes and spermatids and critically regulate their differentiation (Choi et al., 2021; Goh et al., 2015; Wu et al., 2020). Extensive studies have presented compelling evidence supporting different, if not discrepant, conclusions regarding pachytene-piRNA activities, including transposon silencing, genome integrity protection, endogenous mRNA degradation through deadenylation or cleavage, translational activation, and MIWI degradation through ubiquitination (Dai et al., 2019; Goh et al., 2015; Gou et al., 2014; Reuter et al., 2011; Zhang et al., 2015; Zhao et al., 2013; Zheng and Wang, 2012). Perhaps Drosophila endo-piRNAs, conferring sequence diversity like pachytene-piRNAs, exhibit multiple activities. Of note, endo-piRNAs were minor constituents of germline piRNAs in whole testis profiling, however, their abundance in late spermatocytes and spermatids might be underrepresented. Further dissection of endo-piRNA activities could reveal the commonalities and differences of piRNA-mediated sperm formation between animals. Hsp90 machinery controls the ligand binding processes of selected client proteins often through dynamic and transient interactions (Schopf et al., 2017). These chaperone functions include loading/binding of small RNA precursors (ligands) to Ago family proteins (clients). Once loading occurs, Hsp90 changes its dimer conformation through ATP hydrolysis, and dissociates from Ago complexes to complete RISC formation (Iki et al., 2010; Iwakawa and Tomari, 2022; Iwasaki et al., 2010, 2015; Miyoshi et al., 2010; Tsuboyama et al., 2018). Cochaperone Cyp40 functions with Hsp90 during RISC formation, and transiently interacts with Ago proteins in cell lysate systems (Iki et al., 2012, 2020). However, the transient-thus-unstable binding nature makes cochaperone-client interactome difficult in tissue levels. By overcoming the above challenges with the proximity identification method, BioID/TurboID (Branon et al., 2018; Roux et al., 2012), this study identified otherwise undetected physical and functional links between Cyp40 and selected Ago members in testicular germ cells. BioID/TurboID combined with different genetic tools available in model organisms will further decipher selective (co)chaperone-client interactions within and beyond our scopes.

Lastly, the fact that endogenous genes can produce both proteins and regulatory RNAs raises a question of how sequences coding multiple functional factors arrange their genetic information through evolution. It would be interesting to investigate how protein-coding genes evolve in organisms utilizing endo-piRNAs compared to those lacking the biogenesis, and if there are any evidence supporting the existence of selective pressures directing functionalization of endo-piRNAs over proteins.

## Materials and Methods

### Fly stocks and cultures

Fly stocks are reared at room temperature (RT) on a molasses/yeast medium (5%(w/v) dry yeast, 5%(w/v) corn flower, 2%(w/v) rice bran, 10%(w/v) glucose, 0.7%(w/v) agar, 0.2%(v/v) propionic acid, 0.05%(v/v) p-hydroxy butyl benzoic acid). The following stocks were used: *eGFP-Piwi (attP2)* (Sienski et al., 2012), e*GFP-Aub/CyO* [DGRC#118621] (Kina et al., 2019), *cyp40*^*KO*^/*TM3* (Iki et al., 2020), Df(3L)BSC669/*TM6C* [BL#26521] (deficiency for *cyp40*), *bam-Gal4/TM6, FLAG-HA-ago2* (Czech et al., 2008a), *yellow white (y w), ago2*^*454*^/*TM3* [BL#36512], *Df(3L)BSC558/TM6C* [BL#25120] (deficiency for *ago2*), *dcr2*^*L811fsX*^ [BL#33053], *aub*^*HN2*^/*CyO* [BL#8517], *aub*^*N11*^/*CyO* (Harris and Macdonald, 2001), *ago3*^*t2*^/*TM6B* [BL#28269], *ago3*^*t3*^/*TM6B* [BL#28270], *rhi*^*02086*^/*CyO* [BL#12226], *Df(2R)Exel7149/CyO* [BL#7890] (deficiency for *rhi*), *protamine-B-eGFP*/*CyO* [DGRC#109173], *nos-phiC31*; P{y[+t7.7]=CaryP}attP40 [BL#25709], *nos-phiC31*;; PBac{y[+]-attP-3B}VK00033 [BL#32543].

### Plasmid construction and fly transformation

All the primers used for plasmid construction are listed in Table S4. To generate UASp-*mTurbo-FLAG-cyp40* or UASp-*mTurbo-FLAG-cyp40*^*RKAA*^, *mTurbo* was amplified by pfusion PCR (NEB) using Ti833/Ti834 as primers and 3xHA-miniTurboNLS_pCDNA3 (addgene #107172) as template. *FLAG-cyp40* or *FLAG-cyp40*^*RKAA*^ fragment was amplified by PCR using primers Ti835/Ti836 and UASp-*GFP-FLAG-cyp40* or UASp-*GFP-FLAG-cyp40*^*RKAA*^ as template (Iki et al., 2020). Amplified *mTurbo* and *FLAG-cyp40* fragments were introduced into XbaI site of pUASp-K10-attB vector using In-Fusion HD Cloning Kit (Takara). To generate UASp-*mTurbo-FLAG-cyp40*^*deltaTPR*^, a fragment containing *mTurbo-FLAG-cyp40*^*deltaTPR*^ (primers: Ti835/484) was amplified by PCR using UASp-*mTurbo-FLAG-cyp40* as template, and introduced into XbaI site of pUASp-K10-attB vector. The constructs were integrated into *attP40* site. To generate UASp-*mTurbo-FLAG-aub* (only used for rescue experiment in this study), *FLAG-aub* fragment (primers: Ti953/894) was amplified by PCR using *y w* cDNA as template. *mTurbo* and *FLAG-aub* fragments were introduced into XbaI site of pUASp-K10-attB vector. The construct was integrated into *attP-3B* site.

### Proximity-dependent biotin identification in testicular germ cells

mTurbo fusion proteins were expressed in germline-restricted manner under *bam* promoter activity using Gal4/UASp system (Rørth, 1998). After eclosion, male progenies were reared at 25°C for three days in the modified molasses/yeast medium supplemented with 100μM biotin (Nacalai). From two hundred testes, biotinylated proteins were purified as described in (Roux et al., 2012) with some modifications. After exchanging the buffer with 100μl PI-lysis buffer (50mM Tris-HCl [pH7.5], 500mM NaCl, 2mM EDTA, 2mM DTT, 0.4%[w/v] SDS, cOmplete Protease Inhibitor Cocktail Tablet [Roche]), testes were homogenized using Bioruptor (Diagenode) for 30sec (power H) for six times with 30sec intervals. Triton-X100 was added to sample mixtures at a final concentration of 2%[v/v], and homogenization was further performed for 30sec for three times with 30sec intervals. After centrifugation (20,000×g, 10min, 4°C), supernatant was diluted by mixing with equal amount of 50mM Tris-HCl [pH7.5] solution, and incubated with pre-equilibrated 15μl slurry volume of Dynabeads MyOne Streptavidin C1 (Thermo) overnight at 4°C with gentle rotation. Next day, beads were washed twice with W1-buffer (50mM Tris-HCl [pH7.5], 250mM NaCl, 0.2%[w/v] SDS, 1mM EDTA, 1mM DTT), twice with W2-buffer (50mM HEPES-KOH [pH7.4], 500mM NaCl, 0.1%[w/v] deoxycholate [Nacalai], 1%[v/v] Triton-X100, 1mM EDTA), twice with W3-buffer (10mM Tris-HCl [pH8.0], 250mM LiCl, 0.5%[w/v] deoxycholate, 0.5%[v/v] NP-40 (Nacalai), 1mM EDTA), and twice with W4-buffer (50mM Tris-HCl [pH7.5], 50mM NaCl). Proteins bound to magnet beads were denatured by boiling at 95°C for 5min in 2x protein loading buffer (4%[w/v] SDS, 200mM DTT, 0.1%[v/v] BPB, 20%[v/v] glycerol) saturated with biotin, resolved by SDS-PAGE in 5-20% precast gel (ATTO). After electrophoresis, proteins were visualized by using Silver Stain MS Kit (WAKO). Proteins in gel particles were digested with trypsin and analyzed by LC-MSMS using Q-Exactive (Thermo) and UltiMate 3000 Nano LC (Thermo) in CoMiT Omics Center (Osaka Univ, Japan). The mass spectrum data were analyzed by Mascot v2.5.1 (Matrix Science). Proteins were identified with threshold of 1 peptide minimal. Pseudo value 0.5 was added to all identified proteins, then peptide count values were normalized by using total peptide counts in a certain condition.

### Small RNA extraction from testes

RNA was extracted from six hundred testes of ≤3 days old using miRNeasy (QIAGEN). Less than 200 nt fragments were collected using miRelute column (QIAGEN). 2pmol of oligo DNA (5’-AGTCTTACAACCCTCAACCATATGTAGTCCAAGCAGCACT-3’) containing complementary sequence to 2S rRNA was added to 1μg of RNA solution, and 2S rRNA hybridizing with oligo DNA was digested with RNase H (NEB). RNA mixture depleted of 2S rRNA was loaded onto 8M urea-polyacrylamide gel (12%) and size-separated in parallel with RNA ladder (Dynamarker DM253, BioDynamics Laboratory Inc.) in 0.5x Tris-borate EDTA (TBE) buffer. Gel area within the range of 20-30 nt was excised, and contained RNAs were eluted in 300 mM NaOAc (pH 5.3) solution by dilution overnight at 4°C with gentle rotation. RNA fragments were precipitated in the presence of 80% (v/v) ethanol and 40μg/ml glycogen (Nacalai). RNA pellet was rinsed twice with 80%(v/v) ethanol, then resuspended in RNase-free water.

### Piwi or Aub immunoprecipitation and bound piRNA extraction

Five hundred testes of ≤3 days old were homogenized using plastic pestle (Biomasher, Nippi) in IP-NX buffer (30mM HEPES-KOH [pH7.4], 300mM NaCl, 2mM MgCl_2_, 2mM DTT, 10%[v/v] glycerol, 0.5%[v/v] Triton-X100, complete tablet). After centrifugation (12,000×g, 5min, 4°C), to purify endogenous Aub proteins, the supernatant was incubated with anti-Aub antibodies (1:10) for 3h on ice, then further incubated with the mixture of protein A Dynabeads and protein G Dynabeads (Thermo Fisher Scientific) overnight with gentle rotation. To purify GFP-Aub or GFP-Piwi, the supernatant was incubated with anti-GFP antibody-conjugated magnet beads (MBL) overnight with gentle rotation. The beads were washed 5 times with wash buffer (30mM HEPES-KOH [pH7.4], 800mM NaCl, 2mM MgCl_2_, 2mM DTT, 10%[v/v] glycerol, 0.5%[v/v] Triton-X100, complete tablet). The beads-bound sRNAs were extracted with TRIzol LS reagent (Thermo) following the manufacture’s protocol, and precipitated in the presence of 50%(v/v) isopropanol and 20μg of glycogen (Nacalai) overnight at -20°C. After centrifugation (20,000×g, 20min, 4°C), the pellet was rinsed twice with 80%(v/v) ethanol, then resuspended in RNase-free water.

### Deep-sequencing of small RNAs and data processing

The sRNA-seq libraries were constructed with NEB Next Multiplex Small RNA Library Prep Set for Illumina (NEB). After agarose gel electrophoresis, the fragments containing sRNAs were extracted and sequenced by Illumina HiSeq2500. The library construction and deep-seq were performed in Research Institute for Microbial Disease (RIMD, Japan). Data processing was performed by using CLC Genomics Workbench (QIAGEN). Following the removal of 3’ adaptor sequence (5’- AGATCGGAAGAGCACACGTCT-3’), reads ranging from 23 to 29 nt were analyzed unless otherwise indicated. rRNA-, tRNA-, miRNA-, snRNA-, and snoRNA-mapping reads were considered as non-piRNAs and excluded. Remaining reads were mapped without mismatch to the major piRNA clusters (Chen et al., 2021a). Remaining reads were mapped without mismatch to transposons (dmel-all-transposons-r6.39, flybase). Non-specifically matching reads were mapped randomly. After removing cluster- or TE-mapping piRNAs, the remaining reads were mapped without mismatch to genome (ftp.ensembl.org/pub/current_gtf/drosophila_melanogaster/Drosophila_melanogaster.BDGP6.91.chr.gtf.gz). Maximal number of hits for a read was 1 (This setting includes multi-mappers on a certain gene, but excludes multi-mappers on multiple genes). To construct bedgraph, binary alignment map (bam) data exported from CLC were processed using bedtools (genomecov) with scaling factor calculated by reads remaining after non-piRNA removal. Within corresponding conditions, individual bedgraphs were merged (unionbedg), and then visualized in IGV (https://software.broadinstitute.org/software/igv/). To profile the lengths of gene mapping reads, total RNA bam data were first merged within in-house or public data set (2 or 4 data, samtools merge). In each data set, reads mapping to a certain gene were extracted (samtools view), read length was obtained (samtools stats), the proportion was calculated within 18∼29-nt RNAs, and mean values in 2 data sets were finally obtained. In the case of analyzing gene groups, mean of included genes were further calculated. To analyze nucleotide preference, 6 bam data were merged first and included reads were analyzed altogether for total RNAs, while 2 bam data were merged for Aub-bound RNAs. From merged bam, read sequence information was extracted (BBMap reformat.sh, .bam-to-.fa conversion). 1^st^ to 23^rd^ sequences of individual reads were further extracted (Seqkit), and nucleotide probability was obtained for each nucleotide position using Geom_logo under R environment. To measure the density of Aub-bound piRNAs on 5’UTR, CDS, 3’UTR, reads mapping to respective regions were counted (samtools view). Density was given by dividing read counts with length of 5’UTR, CDS, or 3’UTR. Relative density values were calculated, and mean of Aub-IP and GFP-Aub-IP data were obtained. Values from 14 representative host genes were analyzed by box plot under R (ggplot2). To characterize piRNA strand orientation, mapping was performed in strand-specific manner using CLC (Strand specific setting: Forward or Reverse), and the proportion of sense strand reads in total mapped reads were calculated. To identify 5’-to-5’ complementary 10-nt overlap pairs between siRNAs and piRNAs, or between miRNAs and piRNAs, we developed a software available in https://github.com/Dme-Research/complementary-pair#complementary-pair. The partners of representative hp-derived siRNAs and individual miRNA strands were searched within Aub-bound piRNAs mapping to endogenous gene exons. For this purpose, reads were merged between Aub-IP and GFP-Aub-IP (*cyp40*^*KO/TM6*^ background), or between 2 replicates of Aub-IP (*y w* background). When >10 reads of piRNAs were paired with a certain si/miRNA (software version1_1 requesting 10nt perfect complementarity without GU wobble), piRNA host genes were identified. piRNAs paired to miRNAs were analyzed only when the miRNA strand is detectable in testes and shows sorting preference to Ago2 (RPM Ago1-IP < Ago2-IP) (Iki et al., 2020). To identify endo-piRNAs that can target *nej* and *ssrp* transcripts, endo-piRNAs derived from 14 representative host genes were extracted and mapped (bowtie, up to 3 mismatches allowed from 1^st^ to 15^th^ nucleotides of piRNAs. 15-nt sequences were extracted using Seqkit before mapping). Mapped endo-piRNAs were visualized using IGV, and further filtered through a targeting rule not allowing any mismatch from 2^nd^ to 7^th^ nucleotides of endo-piRNAs (GU wobble allowed). Potential target sites were chosen when major form of targeting endo-piRNAs counted >10. For individual endo-piRNAs predicted to target *nej*, 5’-to-5’ complementary 10-nt overlaps were searched within degradome products or Aub-bound piRNAs derived from *nej*.

### Transcriptome analysis

Two fly lines, {*aub*^*HN2*^*/CyO; ago2*^*454*^*/TM6*} and {*aub*^*N11*^*/CyO; ago2*^*Df*^*/TM6*}, were crossed to obtain male progenies of all genotypes analyzed by transcriptome. From 150∼200 testes of ≤3 days old adults, total RNAs were extracted using TRIzol LS reagent (Thermo) following the manufacture’s protocol, and precipitated in the presence of 50%(v/v) isopropanol and 20μg of glycogen (Nacalai) overnight at -20°C. After centrifugation (20,000×g, 20min, 4°C), the pellet was rinsed twice with 80%(v/v) ethanol, then resuspended in RNase-free water. After treating with DNase I (NEB), RNA mixture was added with equal volume of phenol:chloroform:isoamyl alcohol (25:24:1, pH5.2, Nacalai), vortexed, centrifuged (20,000×g, 3min), and the supernatant was collected. After repeating this step, RNA in the supernatant was precipitated in the presence of 0.3M NaOAc (pH5.2) and 70%(v/v) ethanol overnight at -20°C. After centrifugation (20,000×g, 20min, 4°C), the pellet was rinsed twice with 80%(v/v) ethanol, resuspended in RNase-free water, and stored at -80°C until shipment. Library construction and deep-sequencing were performed by Rhelixa (Japan). Polyadenylated RNA selection was performed with NEBNext Poly(A) mRNA Magnetic Isolation Module. Libraries were constructed with NEBNext Ultra II Directional RNA Library Prep Kit, and sequenced by using NovaSeq 6000 (Illumina). The reads were processed and analyzed with CLC genomics workbench (QIAGEN). Paired reads were mapped in the default setting on *D. melanogaster* genome (reference sequence: BDGP6, ensembl.org). Maximal number of hits for a read was 10. Using read counts on exons from biological duplicate samples, differential expression analysis was conducted with EdgeR under R environment. Differentially expressed genes were selected with the threshold of FDR<0.0001 (intersect), and venn diagram was drawn under R. Genes included in individual categories were listed in Table S3. Gene ontology analyses were performed in WebGestalt (http://www.webgestalt.org/). bedgraph was made by using bedtools and processing bam data exported from CLC. Scaling factor was given by total counts of exon mapped reads. bedgraph was visualized in IGV. For transcriptome data used in Figure 1, RNA (>200nt) was extracted from 400 testes of ≤3 days old flies using miRNeasy Mini column (QIAGEN). For the first replicate, extracted RNA was treated with DNase I (NEB) and Ribo-Zero rRNA removal kit (Illumina). After purifying RNA, the libraries were constructed with TruSeq standard mRNA Library Prep Kit (Illumina), and sequenced with HiSeq3000 (Illumina) in RIMD (Japan). For the second replicate, polyadenylated RNAs were collected after DNase I-treatment. polyadenylated RNA enrichment, library construction (NEBNext Ultra Directional RNA Library Prep Kit for Illumina), and sequencing with HiSeq X (Illumina) were performed by AnnoRoad Co.,Ltd (China).

### Data and Code Availability

All deep-sequencing data analyzed in this study are listed in Table S5. Newly generated data are available with BioProject accession ID: PRJNA870252 (https://dataview.ncbi.nlm.nih.gov/object/PRJNA870252?reviewer=49bs22k4tkhsj1h25upsfeomvn)

### Immunoblotting

Protein samples were denatured by boiling at 95°C for 3min in protein loading buffer (2%[w/v] SDS, 100mM DTT, 0.05%[v/v] BPB, 10%[v/v] glycerol), resolved by SDS-PAGE, and transferred to 0.2*μ*m PVDF membrane (Wako) using the semi-dry system (Trans-blot Turbo, Bio-Rad). The membrane was blocked in 4%(w/v) skim milk (Nacalai) in ×1 phosphate-buffered saline (PBS) supplemented with 0.1%(v/v) Tween-20, and further incubated with the following antibodies; anti-Aub antibody (guinea pig, 1:500) (Lim et al., 2022), anti-GFP antibody (Clontech, rabbit, 1:2000), anti-Ago1 antibody (Abcam, Ab5070, rabbit, 1:1000), anti-Ago2 antibody (guinea pig, 1:100) (Iki et al., 2020), anti-Piwi (mouse, P4D2, 1:10), anti-FLAG M2-peroxidase (HRP) (SIGMA, #A8592, 1:5000), anti-H3K18Ac (active motif, #39756, 1:2000), anti-H3K27Ac (active motif, #39136, 1:2000), anti-H4K8Ac (active motif, #61104, 1:2000), anti-H4K12Ac (active motif, #39928, 1:2000), anti-guinea pig immunoglobulins-HRP (DAKO, 1:1000), anti-rabbit IgG-HRP (BioRad, 1:3000), anti-mouse IgG-HRP (BioRad, 1:3000). The chemiluminescent signals were obtained by using Chemi-Lumi One (Nacalai), and detected by Chemidoc MP Imaging system (BioRad). The images were processed by using Pixelmator.

### Histochemistry and image acquisition

Testes were dissected from adult males in ×1 PBS buffer supplemented with 0.4%(w/v) bovine serum albumin (BSA, Wako), and fixed in 5.3%(v/v) paraformaldehyde (Nacalai) in ×0.67 PBS buffer for 10 min. To observe DNA and individualization complexes, testes were incubated with 1μM 4’,6-Diamidine-2’-phenylindole dihydrochloride (DAPI) and phalloidin Rhodamine X Conjugated (Wako, 1:1000) in PBX buffer (×1 PBS containing 0.2%[v/v] Triton-X100). To observe ProtB-GFP, the emission signal of GFP were acquired. For immunostaining, fixed testes were washed with PBX and incubated with PBX containing 2%(w/v) BSA for 30 min for blocking. The primary antibody incubation was performed overnight at 4 °C, and testes were washed with PBX at 25 °C for 1h. The secondary antibody incubation was then performed at 25 °C for 2h, and then testes were washed with PBX at 25 °C for 1h. The following antibodies were used; anti-Vasa antibody (guinea pig, 1:5000), anti-guinea pig IgG-Alexa Fluor 555 (Molecular probes, 1:500), anti-H3K18Ac (active motif, #39756, 1:200), anti-H3K27Ac (active motif, #39136, 1:200), anti-H4K5Ac (active motif, #39170, 1:200), anti-H4K8Ac (active motif, #61104), anti-H4K12Ac (active motif, #39928, 1:200), anti-H4 (active motif, #61300, 1:200), anti-rabbit IgG-Alexa Fluor 555 (Molecular probes, 1:500). Images were acquired using confocal microscope LSM900 (Zeiss). Images were processing by using Zen (Zeiss) and Pixelmator.

### Quantivative reverse transcription PCR (qPCR)

RNA was extracted from ∼20 testes of ≤3 days old flies for each condition using TRIzol LS reagent (Thermo) following the manufacture’s protocol. Using DNase I (NEB)-treated RNA, cDNA was synthesized with 2.5μM oligo dT adaptor using SuperScript III reverse transcriptase (Thermo). qPCR reaction was performed using KAPA SYBR FAST qPCR Master Mix (Roche) and gene specific primers (Table S4) in Quantstudio5 Real-Time PCR system (ABI).

### Male fertility test

Single male was mated with 6 *y w* virgin females for the first 3 days, and with another 6 *y w* virgin females the following 2 days at 25°C. The total number of hatched eggs was counted.

## Acknowledgement

We thank Miwako Okamura, Chisato Yanagisawa, and Rizky Mutiara for their help with maintenance of fly lines. We are grateful to Bloomington Drosophila Stock Center and Kyoto Stock Center for providing fly stocks. We thank Daisuke Motooka for sRNA-seq services, and Mikiko Siomi for reagents. We also thank all members in our lab for their insightful discussion and suggestions. This work was supported by Grant-in-Aid for Scientific Research C (22K06081) for T.I., and TAKEDA Bioscience Research Grant (J191503009) for T.K.

## Author contributions

T.I. designed the research, performed experiments, and analyzed the data. T.I. performed bioinformatic analyses with S.K. T.I. wrote the manuscript with T.K.

## Competing interest declaration

The authors declare there is no competing interest.

## Notes

### Competing Interest Statement

The authors have declared no competing interest.

