## Supplementary figures and images for "A pathway to produce non-coding piRNAs from endogenous protein-coding regions supports Drosophila spermatogenesis"

### Figure S1

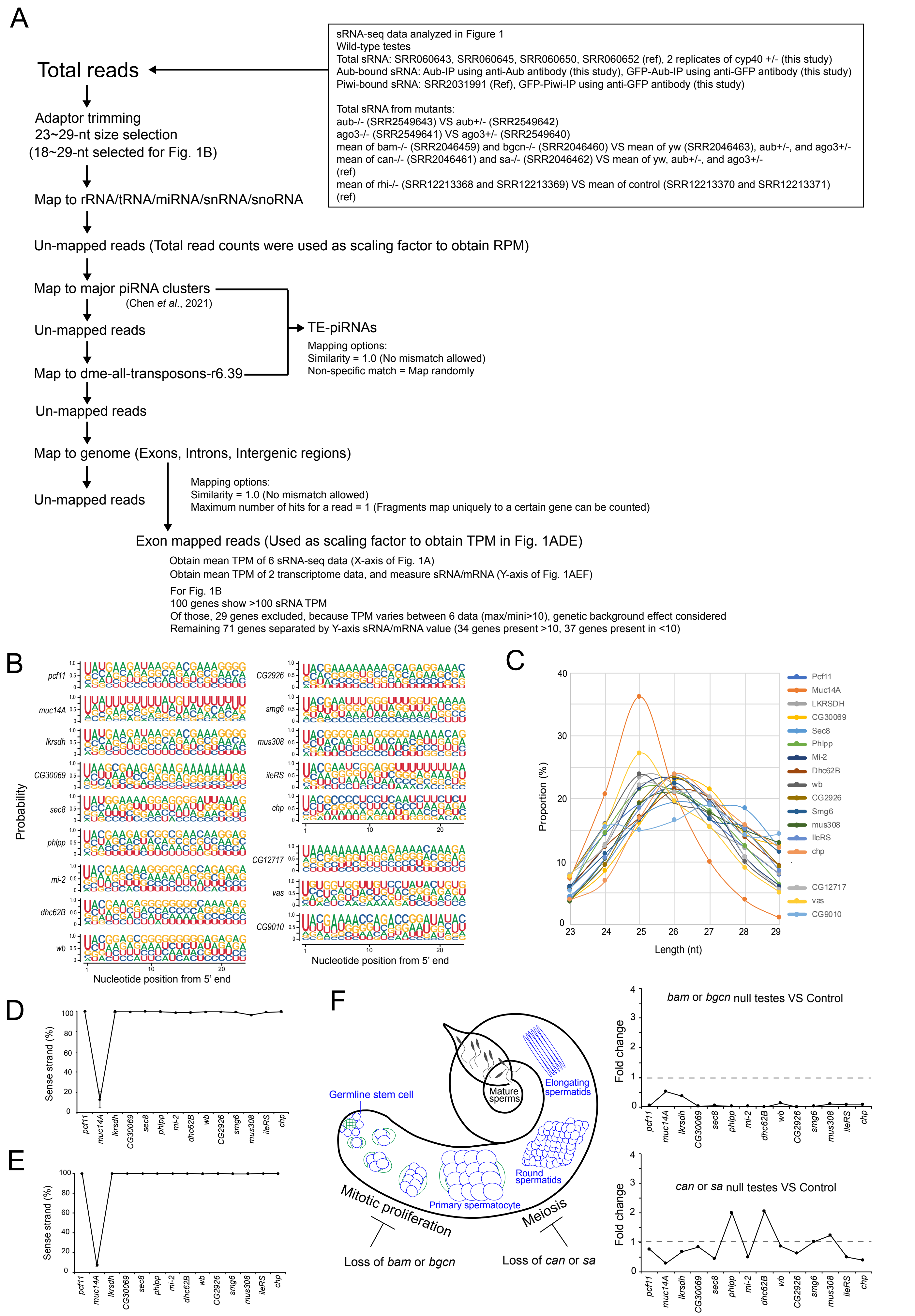

### Figure S4

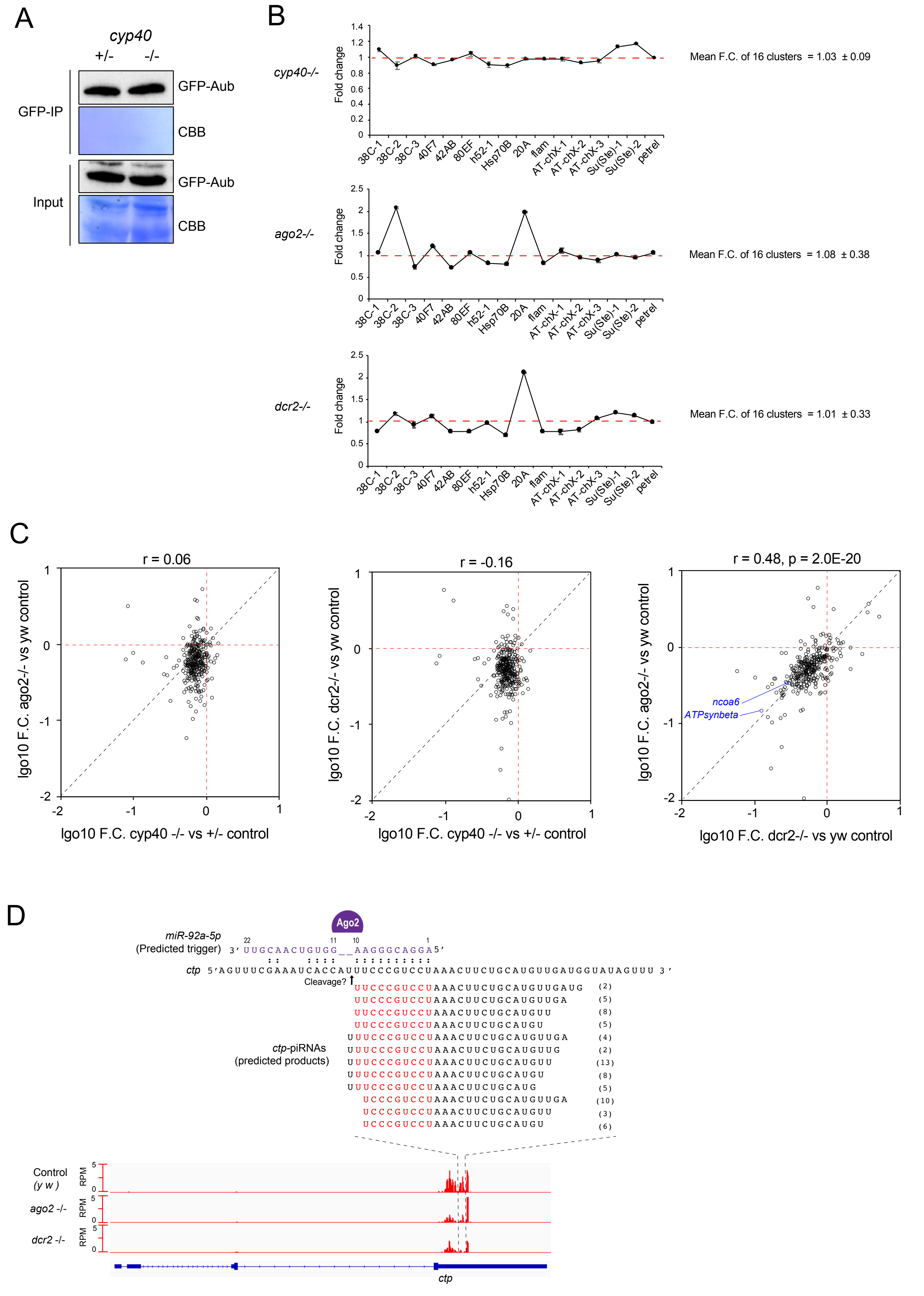

### Figure S5

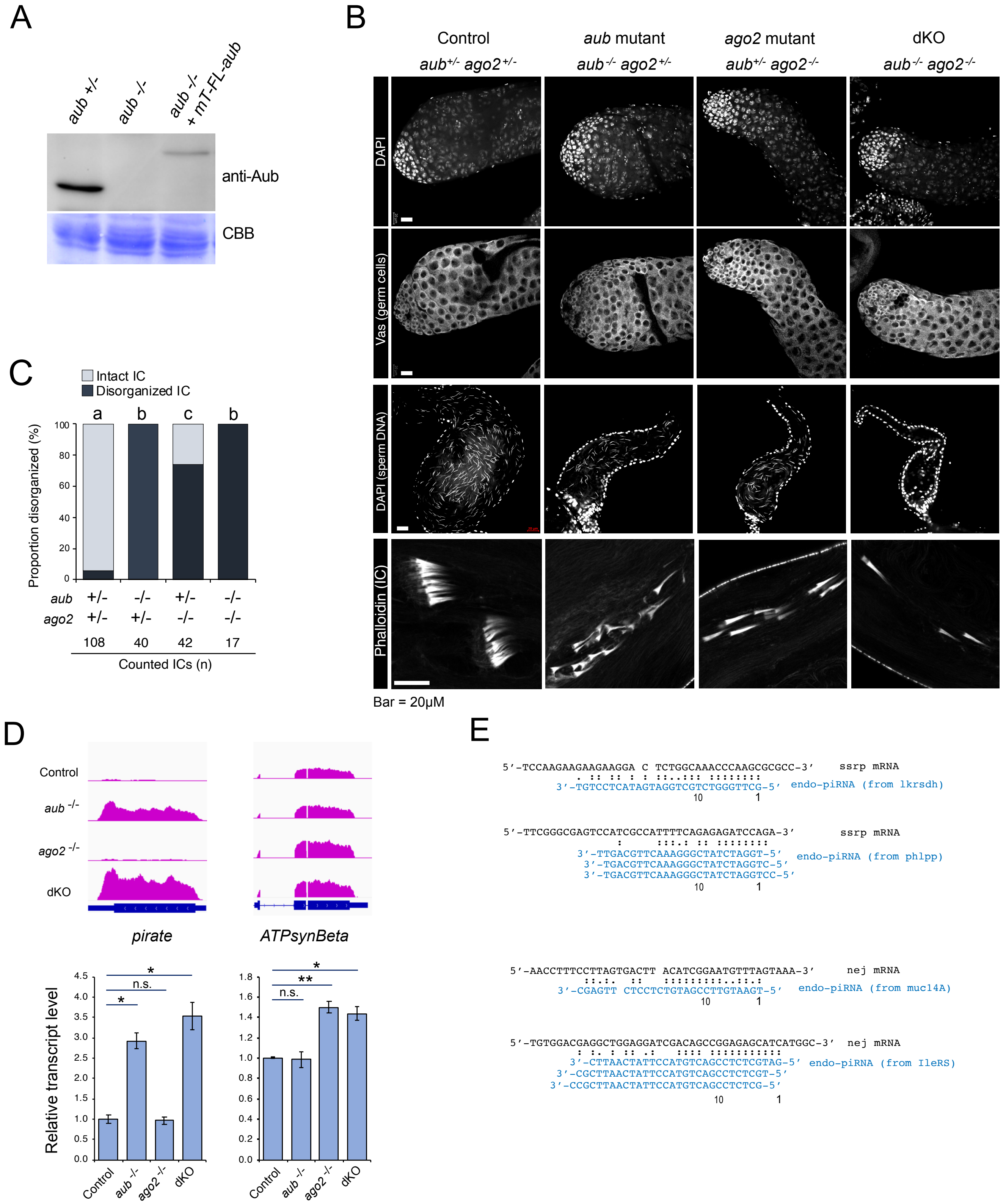

### Figure S6

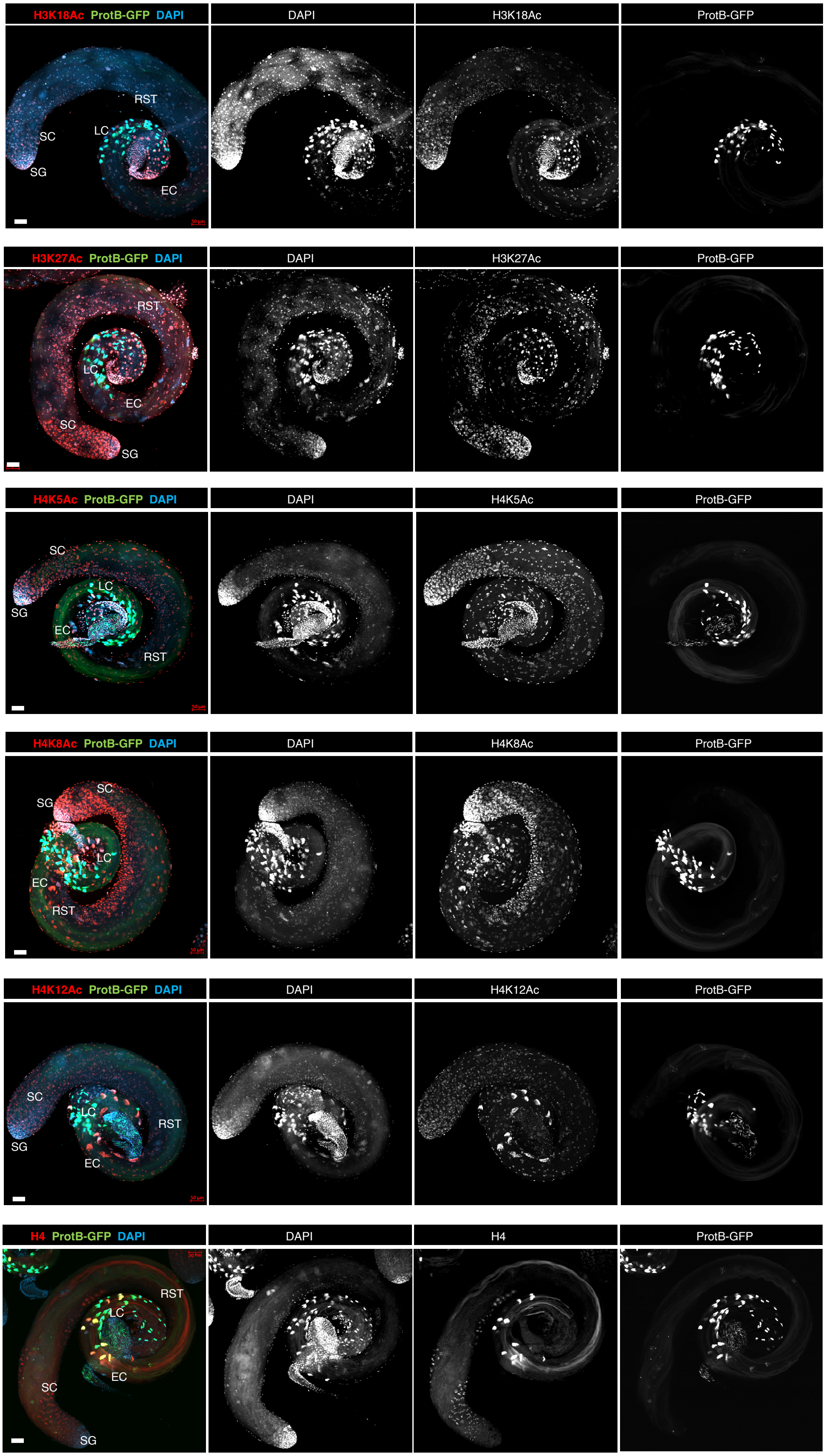
