## Supplementary material for "A pathway to produce non-coding piRNAs from endogenous protein-coding regions supports Drosophila spermatogenesis": Figure S2

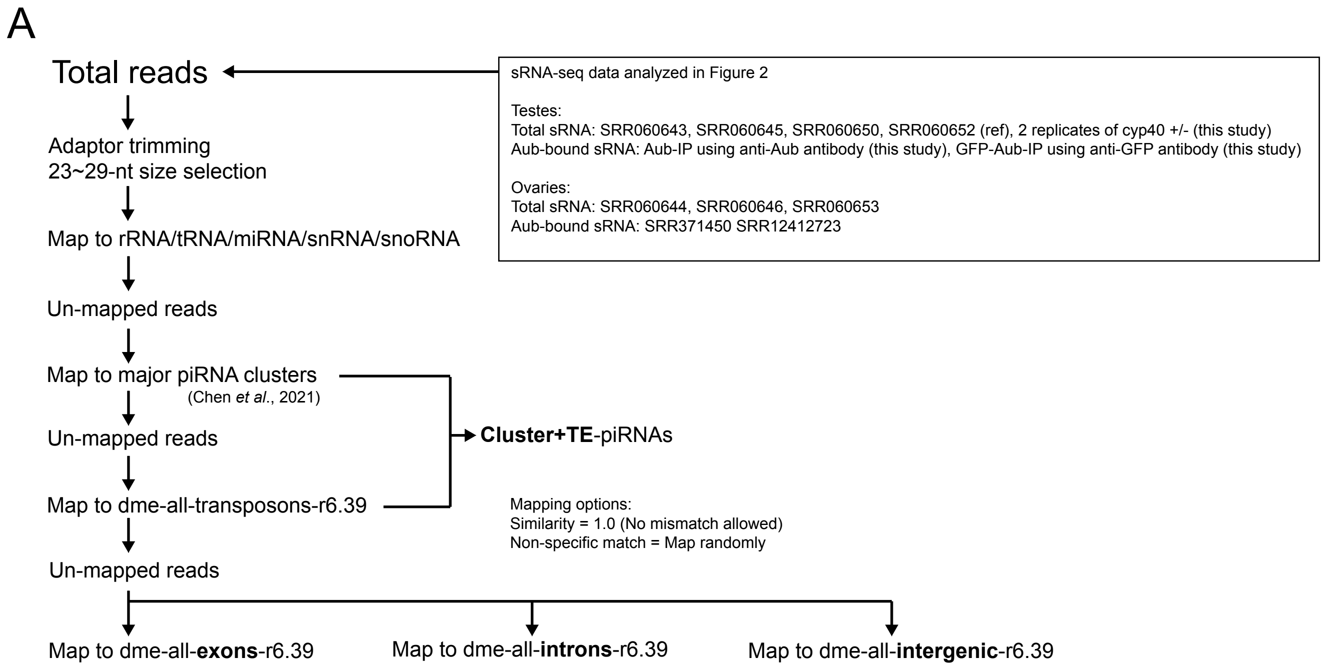


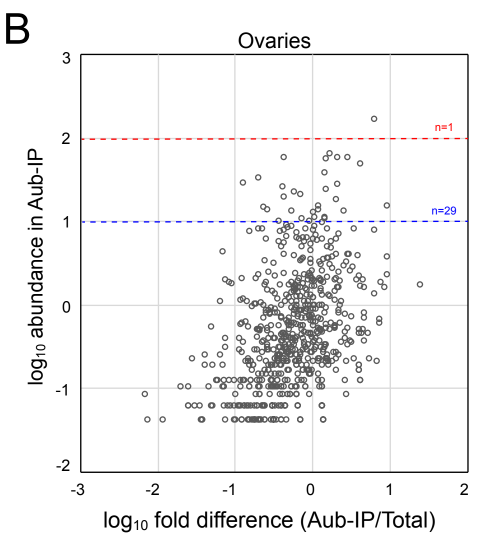


1st filter: 23~29-nt sRNA/mRNA (TPM/TPM) >10
Testis: 2449 genes
Ovaries: 910 genes

2nd filter (loose): Aub-bound piRNA (RPM) >10
Testis: 324 genes
Ovary: 29 genes

2nd filter (strict): Aub-bound piRNA (RPM) >100
Testis: 17 genes
Ovary: 1 gene

C

| Genes | Mean RPM in ovarian Aub-IP |
|---|---|
| RpL10Ab | 162.3244626 |
| RpL3 | 61.36694079 |
| Stt3A | 56.61918361 |
| rl | 56.05558953 |
| CG17514 | 55.86731562 |
| Maf1 | 54.68528491 |
| CG17493 | 47.91498149 |
| Muc14A | 36.34839476 |
| Nipped-A | 32.28901014 |
| CG32000/anne | 27.48714437 |
| CG30069 | 26.58828949 |
| CG17715 | 24.93818175 |
| Pcf11 | 23.49794121 |
| CG40160 | 21.46545766 |
| RpS2 | 20.07012238 |
| sta | 18.62288824 |
| UQCR-11 | 17.37135061 |
| CG11023 | 14.72711646 |
| Sox102F | 14.67000212 |
| Cp15 | 14.19609711 |
| CG41099 | 13.44944461 |
| CG3704 | 13.15681991 |
| l(3)76BDm | 12.99376016 |
| RpL24 | 12.81971897 |
| Atf6 | 12.27502 |
| CG3812 | 10.8319572 |
| toy | 10.59736691 |
| l(2)k0120 | 10.39559687 |
| RpS15 | 10.35786869 |

These genes were identified as piRNA-hosting genes in testes.

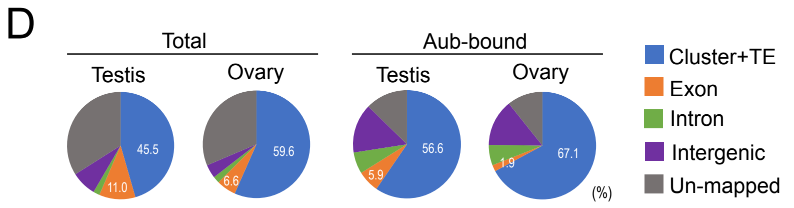


11.0±5.7 6.6±2.0 5.9±1.9 1.9±0.5 Mean proportion of exon-r6.39 mappers

$p$ = 0.18 $p$ = 0.29

Figure S2
