## Supplementary material for "A pathway to produce non-coding piRNAs from endogenous protein-coding regions supports Drosophila spermatogenesis": Figure S3

A

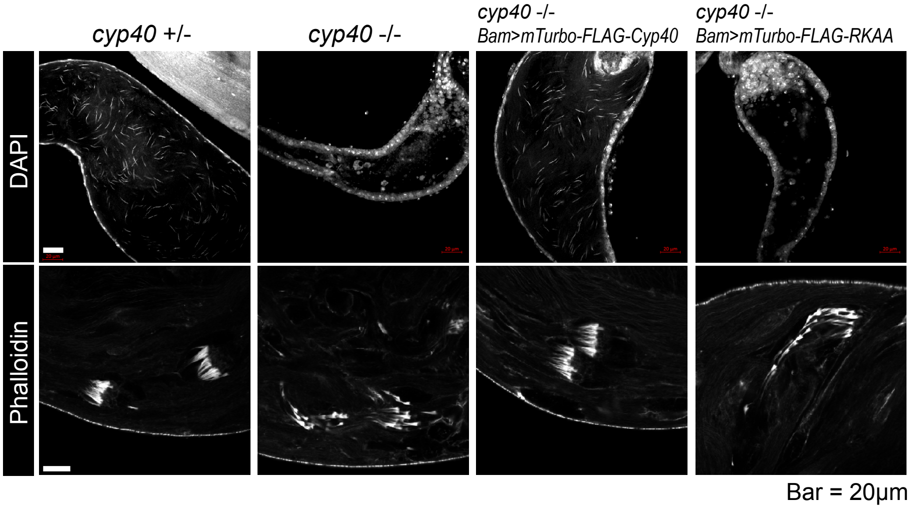


B

| Name | Enrichment | $p$ (<0.01) | Coverage (%) | | | | | |
|---|---|---|---|---|---|---|---|---|
| **Gene silencing** | | **mTurbo:** | **Cyp40** | **Cyp40** | **RKAA** | **ΔTPR** | **GFP** | **GFP** |
| Dcr2 | 12.94 | 2.61169E-06 | 7 | 7 | 0 | 0 | 0 | 0 |
| Ago2 | 11.37 | 0.000122319 | 10 | 8 | 0 | 0 | 0 | 0 |
| Aub | 8.81 | 0.000370819 | 21 | 21 | 1 | 0 | 4 | 0 |
| Qin/Kumo | 5.91 | 0.001117021 | 4 | 4 | 1 | 0 | 0 | 0 |
| fs(1)Yb | 5.29 | 0.00219343 | 5 | 7 | 0 | 0 | 0 | 1 |
| **Nuclear** | | | | | | | | |
| CG9899 | 23.50 | 4.38273E-05 | 51 | 45 | 4 | 0 | 1 | 1 |
| Mle | 12.52 | 0.00015714 | 20 | 24 | 1 | 0 | 3 | 1 |
| Sbf | 10.88 | 0.000216629 | 6 | 8 | 1 | 0 | 0 | 0 |
| Mus308/Pol $\theta$ | 3.02 | 0.008841278 | 1 | 1 | 0 | 0 | 0 | 0 |
| **Microtubule** | | | | | | | | |
| CG17687 | 39.72 | 0.000260524 | 19 | 21 | 0 | 0 | 0 | 0 |
| CG14838/Mmm | 19.74 | 5.19148E-06 | 22 | 19 | 1 | 0 | 0 | 0 |
| Klp10A | 6.06 | 0.000151872 | 7 | 6 | 0 | 0 | 0 | 0 |
| **Mitochondrion** | | | | | | | | |
| Nrd1 | 22.67 | 7.46998E-07 | 20 | 18 | 0 | 0 | 1 | 0 |
| CG3107 | 9.90 | 3.74038E-06 | 7 | 8 | 0 | 0 | 0 | 0 |
| Ogdh | 3.81 | 5.77854E-05 | 2 | 2 | 0 | 0 | 0 | 0 |
| **tRNA metabolism** | | | | | | | | |
| MetRS | 4.92 | 0.005576408 | 40 | 21 | 2 | 3 | 6 | 3 |
| ValRS | 3.81 | 5.77854E-05 | 3 | 2 | 0 | 0 | 0 | 0 |
| **mRNA metabolism** | | | | | | | | |
| Bgcn | 16.88 | 0.004128041 | 8 | 12 | 0 | 0 | 0 | 0 |
| Irp-1a | 12.84 | 0.001757869 | 19 | 11 | 0 | 0 | 0 | 0 |
| Upf1 | 4.59 | 0.003436041 | 2 | 3 | 0 | 0 | 0 | 0 |
| Gld2 | 1.79 | 0.007425165 | 10 | 8 | 3 | 2 | 5 | 4 |
| **Other** | | | | | | | | |
| Ubr1 | 5.98 | 0.004688514 | 2 | 4 | 0 | 0 | 1 | 0 |
| CG14411 | 5.33 | 1.62642E-05 | 6 | 6 | 0 | 0 | 0 | 0 |
| Synj | 3.02 | 0.008841278 | 2 | 1 | 0 | 0 | 0 | 0 |
| Naa15-16 | 3.02 | 0.008841278 | 3 | 1 | 0 | 0 | 0 | 0 |

C

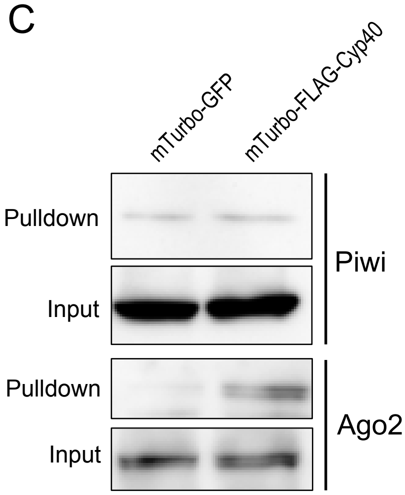


Figure S3
