## Supplementary figure legends for "A pathway to produce non-coding piRNAs from endogenous protein-coding regions supports Drosophila spermatogenesis"

**Figure S1. related to Figure 1**

(A) Scheme for deep-sequencing data processing. (B, C) Nucleotide probability and size distribution of 23~29-nt RNAs in Aub-bound fraction. Reads in 2 data (Aub-IP and GFP-Aub-IP) were merged and analyzed. (D, E) Sense/antisense strand bias of 23~29-nt RNAs mapping to indicated genes. In panel D, in-house and public data for testicular total small RNAs were analyzed separately, and mean values of sense strand proportion were obtained. In panel E, 2 data for Aub-bound RNAs (Aub-IP and GFP-Aub-IP) were analyzed separately, and mean values of sense strand proportion were obtained. (F) Effect of loss of germline differentiation factors on the expression of 23~29-nt RNAs in testes. *bag of marbles (bam)* or *benign gonial cell neoplasm (bgcn)* mutants accumulate early stage spermatogonial cells, while *cannonball (can)* or *spermatocyte arrest (sa)* mutants accumulate primary spermatocytes.

**Figure S2. related to Figure 2**

(A) Scheme for deep-sequencing data processing for comparing testes and ovaries. (B) The pie charts show the proportions of 23~29-nt RNAs mapping to indicated genomic contents. (C) Search for endogenous protein-coding genes accumulating 23~29-nt RNAs in ovaries. As in analyses on testicular RNAs, 910 genes were first extracted by the abundance of 23~29-nt fragments relative to mRNAs. Of those 910 genes, only one gene produced >100 RPM piRNAs, and only 29 genes produced >10 RPM piRNAs in Aub-bound fraction. (D) List of candidate 29 genes producing Aub-interacting piRNAs in ovaries. Genes in red color were identified in testes as hosts of piRNAs. The differences of piRNA abundance between testes and ovaries can be seen in Figure 2B.

**Figure S3. related to Figure 3**

(A) Defective spermatogenesis in testes lacking *cyp40* (-/-; *cyp40^KO/Df^*), and the rescue by *mTurbo-FLAG-cyp40* transgene expressed under germline-specific *bam* promoter activity. Nuclei of mature sperms stored in seminal vesicles were observed with DAPI, and individualization complexes (ICs) in elongating spermatids were observed with phalloidin. *cyp40^RKAA^* (*mTurbo-FLAG-RKAA*) is a non-functional variant of *cyp40*. Heterozygous sibling (+/-) serves as wild-type control. (B) List of proteins identified in the physical proximity of Cyp40. The values for enrichment and statistic difference (*p*) correspond to those shown in Figure 3C. Table also contains peptide coverage (%) of identified proteins. Identified proteins were classified into groups by their known or predicted functions. (C) Immunoblotting of Piwi and Ago2 present in testes (Input) and purified with streptavidin (Pulldown).

**Figure S4. related to Figure 4**

(A) Immunoblotting of GFP-Aub proteins present in testes of heterozygous (+/-) and homozygous (-/-) of *cyp40* (Input) and those purified with anti-GFP antibodies for deep-sequencing (GFP-IP). CBB staining serves as protein loading control. (B) Effect of loss of *cyp40*, *ago2*, or *dcr2* on cluster-mapping piRNA abundance inside Aub-RISCs. Mean ± s.d. of 2 data set was shown. (C) Correlation analyses. F.C. of piRNA abundance was compared between two conditions. Correlation coefficient (r) and p value are shown. (D) 5’-to-5’ complementary overlap screening predicted Ago2-sorted *miR-92a-5p* and *ctp*-derived piRNAs as trigger and product pairs. Note that *ctp*-piRNAs are derived from 3’UTR, as an exception of endo-piRNAs.

**Figure S5. related to Figure 5**

(A) Immunoblotting of Aub proteins in testes of heterozygous control (+/-), or of *aub^N11/HN2^* null mutants (-/-) in the absence or presence of *mTurbo-FLAG*-*aub* transgene expression. CBB staining serves as protein loading control. (B) Phenotypes of testes lacking *aub*, *ago2*, or both. Male progenies of *aub^N11/HN2^* mutant (*aub^-/-^ ago2^+/-^*), *ago2^454/Df^* mutant (*aub^+/-^ ago2^-/-^*), double knock-out (dKO) (*aub^-/-^ ago2^-/^*^-^), or heterozygous sibling control (*aub^+/-^ ago2^+/-^*) were obtained from a single mating. Top two panels show DAPI and Vas (germline marker) signals in apical end of testes containing spermatogonia and spermatocytes. Lower two panels show nuclei of sperms stored in seminal vesicles (DAPI), and ICs formed by spermatids (Phalloidin). (C) Disorganized IC counting. (D) Transcriptome (bedgraph) and qPCR measurement (bar graph) data on *pira* and *ATPsynbeta*. *pira* is regulated by *aub* but not by *ago2*, while *ATPsynbeta* is regulated by *ago2* but not by *aub*. (E) Examples of endo-piRNAs predicted to recognize *ssrp* and *nej*.

**Figure S6. related to Figure 6**

Histone H3 H4 acetylation patterns in testes expressing ProtB-GFP. SG; spermatogonia, SC; spermatocytes, RST; round spermatids, ES; elongating spermatids, EC; early canoe stage spermatids, LC; late canoe stage spermatids, IS; individualizing spermatids. The acetylation patterns were summarized in Figure 6A.
